# Human Myelin Spheres for *In Vitro* Oligodendrocyte Maturation, Myelination and Neurological Disease Modeling

**DOI:** 10.1101/2025.05.28.656435

**Authors:** Karan Ahuja, Roya Ramezankhani, Xinyu Wang, Thibaut Burg, Giulia Amos, Katrien Neyrinck, Alessio Silva, Geethika Arekatla, Eleanor Eva Cassidy, Fatemeharefeh Nami, Joke Terryn, Keimpe Wierda, Katlijn Vints, Niels Vandamme, Suresh Poovathingal, Ivo Lambrichts, Johannes V. Swinnen, Ludo Van Den Bosch, Lies De Groef, Lieve Moons, Catherine Verfaillie, Johan Neyts, Dirk Jochmans, Yoke Chin Chai

## Abstract

Demyelinating diseases, such as multiple sclerosis, damage the protective myelin sheaths of the central nervous system. The development of effective therapies has been hampered by the lack of models that accurately replicate human myelin biology. Here we present a novel method to generate human myelin spheres (MyS) by coculturing of hPSC-derived neuronal and oligodendrocyte precursor cells, to create myelinated neurons. Using multimodal analyses including confocal and (electron)microscopy, single-nuclei transcriptomics, lipidomics, and electrophysiology, we demonstrate myelination in MyS as early as six weeks into coculture. These myelinated structures mature over time into multilamellar and compacted myelin sheaths with lipid compositions and transcriptomic profiles mirror the temporal dynamics of *in vivo* human oligodendrocyte development and neuronal myelination, resembling those of late fetal/postnatal oligodendrocytes. By employing lysolecithin-induced demyelination and Rabies virus infection experiments, we demonstrate the potential of MyS as an innovative, physiologically relevant platform for studying myelin-related neurodegeneration and neuroinfection.

## Introduction

Oligodendrocytes, the myelinating cells of the human central nervous system (CNS), are vital for axonal insulation, efficient action potential conduction, and the overall maintenance of neuronal health. Myelination is a dynamic and lifelong process, which relies on the coordinated interactions between oligodendrocytes and neurons, leading to the synthesis and maintenance of lipid-rich myelin sheaths that are crucial for proper CNS function. Disruption of this process is implicated in a range of demyelinating disorders, such as Multiple Sclerosis (MS), Neuromyelitis Optica, and Leukodystrophies. Recent studies have shown significant species-specific differences in the structure and function of the myelin sheath, particularly between rodents and humans. These differences encompass proteomic^1^ and lipidomic composition^2^, structural organization (such as variations in myelin thickness and internode length) and the timing of oligodendrocyte maturation^3^. Hence, current rodent models fail to capture the intricate processes of human myelin biology, limiting insights into its role in health and disease. Consequently, there is a critical unmet need for a human myelin model that faithfully replicates the temporal maturation stages of myelin sheath formation – from early loose wrapping around axons to the compact, multilamellar myelin characteristic of mature oligodendrocytes. Such a model would offer transformative insights into myelin formation, its disruption in disease and remyelination strategies upon injury, paving the way towards novel therapeutic strategies to treat demyelinating disorders.

Advances in human pluripotent stem cell (hPSC) and brain organoid technologies have led to the development of self-organizing three-dimensional (3D) models that closely recapitulate the neural cytoarchitecture and transcriptional programs of *in vivo* human brain development. However, in most current models, axonal myelination typically does not occur until at least 8-12 weeks in culture^4,5^ (Table S1). These hPSC-derived organoids are usually initiated from a single pluripotent population that undergoes spontaneous or directed differentiation, thereby mimicking, to some extent, early brain developmental trajectories. Like *in vivo* development, oligodendrocyte lineage cells appear later in culture and require additional time to mature. In the human brain, the transition from oligodendrocyte precursor cells (OPCs) to mature, myelinating oligodendrocytes begins in late fetal development and continues well into the postnatal period, especially during the first few years of life^6^. As a result, the existing *in vitro* hPSC models predominantly capture only early-stage neuron-glia interactions, while oligodendrocyte maturation and myelin formation remain largely immature. This immaturity is evidenced by the scarcity of compact, multilamellar myelin sheaths and the presence of immature transcriptional signatures in oligodendrocyte populations^7,8^.

We developed a novel hPSC-derived 3D spheroid model known as Myelin Spheres (MyS), generated by coculturing neural precursor cells (NPCs) with pre-differentiated OPCs to study myelination. As a control, Neuron Monoculture Spheroids (NMS) composed of NPCs alone were also generated. Using a combination of single-nuclei RNA sequencing, lipidomics, electrophysiology and electron microscopy, we demonstrated that MyS supported significantly greater oligodendrocyte maturation and compact myelin formation compared to NMS. These findings demonstrate that incorporating pre-differentiated OPCs accelerates and enhances myelination, offering a more physiologically relevant platform for modeling myelin-related diseases. As a proof-of-concept, MyS was used as a disease modeling platform to study two pathological paradigms: lysolecithin-induced demyelination and Rabies virus infection.

## Results

### 1. Generation and characterization of Neuron Monoculture Spheroids (NMS) and Myelin Spheres (MyS)

We cocultured pre-differentiated cortical neural precursor cells (NPCs) and inducible *SOX10* (i*SOX10*) oligodendrocyte precursor cells (OPCs) using a hanging drop method to generate MyS (Figure 1a). As control, NMS were generated by culturing only NPCs in hanging drops. Both NMS and MyS were cultured in medium supplemented with growth factors supporting oligodendrocyte differentiation for 6-15 weeks. Throughout the whole culture period, NMS were larger in diameter than MyS, likely due to higher initial number of more proliferative NPCs, as compared to MyS (Figure 1b).

**Figure 1:**
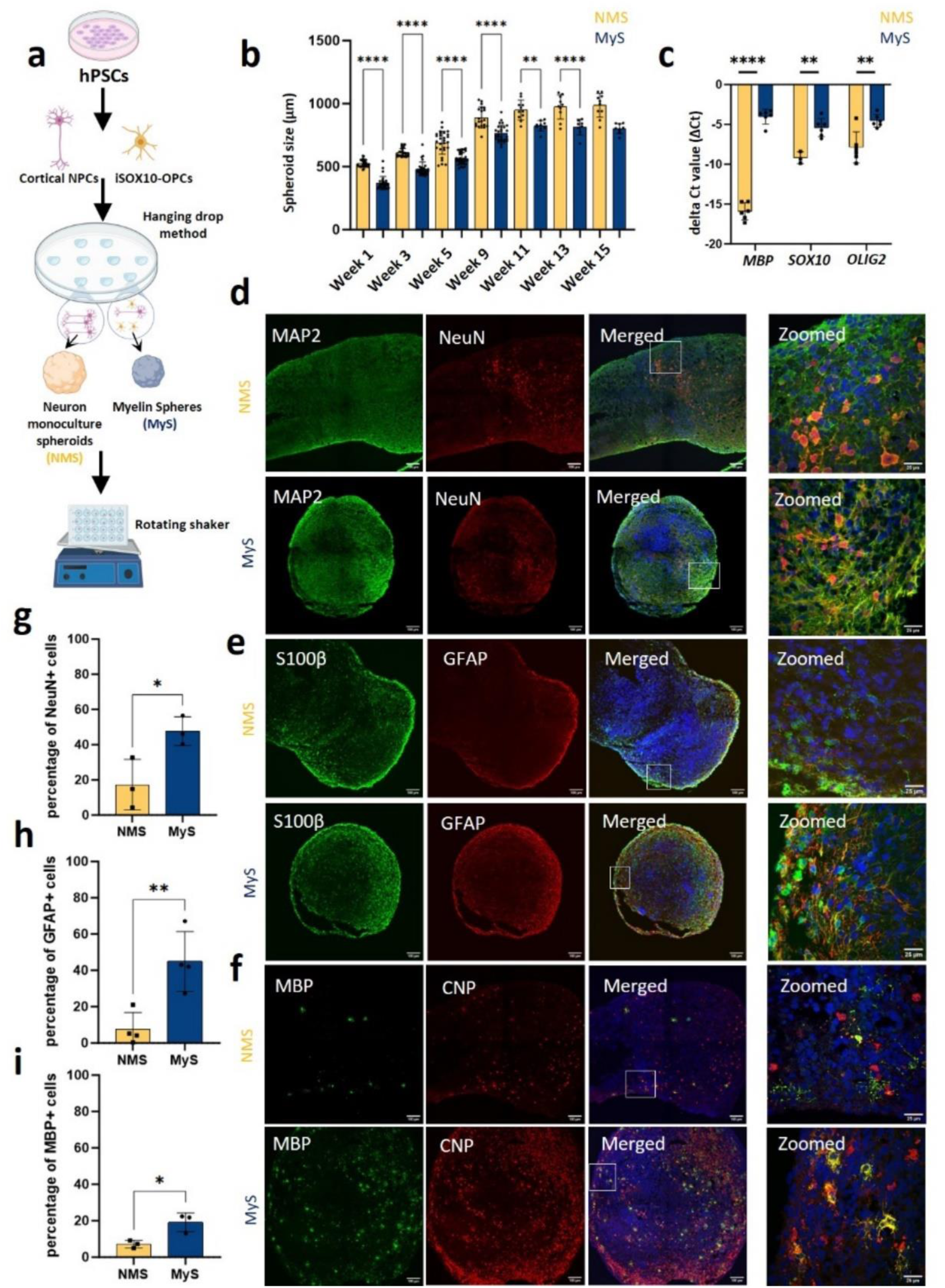
Generation and characterization of Myelin Spheres (MyS) and Neuron Monoculture Spheroids (NMS). **a** Schematic representation depicting the generation of MyS and NMS by coculturing human pluripotent stem cells (hPSCs) derived pre-differentiated cortical neural (NPCs) and oligodendrocyte precursor cells (OPCs) in hanging drops followed by dynamic culture. **b** Growth of Mys and NMS over 15 weeks of dynamic culture indicated by the quantified diameters of MyS and NMS on the brightfield images. **c** Gene expression analysis by RT-qPCR showing significantly higher transcript levels of oligodendrocyte markers (*MBP*, *SOX10* and *OLIG2*) in week 10 MyS as compared to NMS. **d-f** Confocal microscopic images showing the expression of neuronal (MAP2, NeuN), astrocytic (S100β, GFAP) and oligodendrocyte (MBP, CNP) markers in week 10 NMS and MyS via immunostaining (scale bar = 100 μm and 25 μm for zoomed images). Nuclei were counterstained with Hoechst. **g-i** Quantification based on immunostaining images showing significantly higher percentage of NeuN, GFAP and MBP positive cells in week 10 MyS than NMS. Statistical analyses were performed by two-way ANOVA with Sidak’s comparison test (**b**) or unpaired two tailed *t*-tests (**c**, **g-i**). Data shown are mean ± SD. *p< 0.05, **p< 0.01, ****p< 0.0001. N=1 (n=8-32 spheroids, **b**), N=3 (n=3-6 spheroids, **c**), representative immunostaining images for N=3 (**d-f**), N=3 (n=3-4, each data point represents 3 slices, **g-i**).

By real-time quantitative polymerase chain reaction (RT-qPCR) we observed that expression of key transcripts for NPCs (*NESTIN, SOX2*), maturing neurons (*TUBB3, MAP2)* and astroglial (*S100 β, SOX9, AQP4 and ALDH1L1*) in week 10 NMS and MyS were similar (Figure S1a-b). Immunostaining for neuronal (MAP2, NeuN, CTIP2, SATB2) and astrocytic markers (GFAP, EAAT1, S100β) (Figure 1d–e, Figure S1c–d) revealed the presence of well-defined neuronal and astroglial populations. Quantification of NeuN⁺ neurons and GFAP⁺ astrocytes indicated a higher abundance of mature neurons and astrocyte-lineage cells in MyS compared to NMS (Figure 1g– h). We also performed RT-qPCR and immunostaining for OPC- and oligodendrocyte-specific markers on week 10 NMS and MyS. RT-qPCR confirmed the presence of higher levels of OPC (*SOX10, OLIG2*) and oligodendrocyte (*MBP*) transcripts in MyS compared to NMS (Figure 1c). Immunostaining revealed that both NMS and MyS contained cells expressing oligodendrocyte specification transcription factors (OLIG2, SOX10) (Figure S1e) and mature oligodendrocyte markers (MBP, CNP) (Figure 1f). The presence of cells with an OPC or mature oligodendrocyte phenotype in NMS is likely due to the use of oligodendrocyte maturation medium (OMM) containing cytokines that support oligodendrocyte generation and maturation. Nevertheless, a significantly greater percentage of mature oligodendrocytes was present in MyS compared to NMS (Figure 1i).

### 2. MyS-derived oligodendrocytes closely resemble adult human oligodendrocytes

To gain deeper insight into the dynamics of cell lineage progression in MyS and NMS, we performed single-nuclei RNAseq (snRNAseq) at both 8-week and 15-week time points. We first compared the impact of adding i*SOX10*-OPCs to the MyS, but not the NMS on oligodendrocyte development, by combining the 8- and 15-week datasets for each model. By categorizing cells based on their gene expression profiles using Uniform Manifold Approximation and Projection (UMAP) dimensionality reduction (Figure 2a), we identified clusters according to established markers for progenitor cells (*TOP2A* and *MKI67*), oligodendrocytes (*MBP* and *PLP1*), OPCs (*PCDH15* and *PDGFRα*), astrocytes (*GFAP* and *AQP4*), and neurons (*RBFOX3*, *BCL11B*, *SLC17A7, NEUROD6, GAD1* and *GAD2*) (Figure 2b and Figure S2a-d). To confirm the identity of the cellular clusters, we performed gene ontology (GO) enrichment analysis on the differentially expressed genes (DEGs) with higher expression in each cluster. This validated the identities of the clusters previously inferred based on marker expression (Figure 2c and Figure S2e). Notably, the term “Myelination (GO: 0042552)” emerged as the most enriched term in the GO biological processes analysis for the cluster identified as oligodendrocytes (Figure 2c, top). Additionally, the gene expression profile in this cluster significantly matched that of oligodendrocytes in both mouse and human brains, according to the CellMarker 2024 datasets (Figure 2c, bottom).

**Figure 2:**
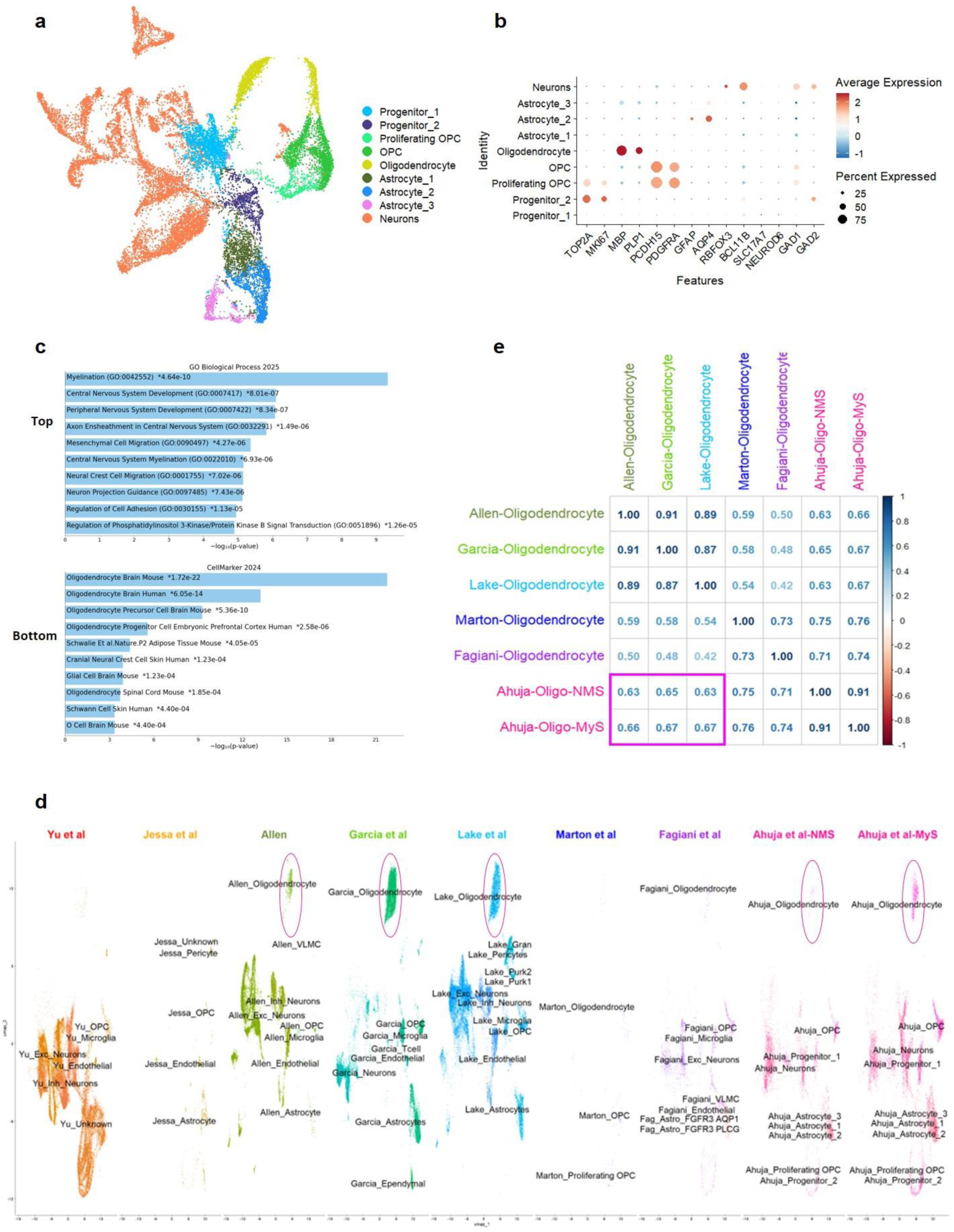
Single-nuclei RNA sequencing results unveiled a notable transcriptomic congruence between mature oligodendrocytes and MyS- or NMS-derived oligodendrocytes. **a** The UMAP embedding of MyS and NMS revealed distinctive clusters characterized by specific transcriptomic signatures of different brain cells. **b** Dotplot shows the average expression and percentage of cells expressing markers for progenitor cells, OPCs, oligodendrocytes, astrocytes, and neurons within each cell cluster. **c** Enrichment analysis of the oligodendrocyte cluster highlighted a significant enrichment of the BP-annotated myelination pathway based on the differentially upregulated genes. Additionally, the cells were matched to those of oligoendrocytes in both mouse and human brains, according to the CellMarker 2024 database (https://maayanlab.cloud/Enrichr/). d. Cells derived from MyS and NMS exhibit transcriptomic profiles more similar to *in vivo* brain cells and other *in vitro* models (the oligodendrocytes are highlighted within the pink ellipse), shown by the UMAP representation of integrated sn/scRNA-seq datasets, which include MyS and NMS samples, *in vivo* adult and developing brain cells, and two *in vitro* brain models. **e** Oligodendrocytes present in MyS and NMS are more similar to adult oligodendrocytes from three different *in vivo* datasets^20–22^ compared to oligodendrocytes developed in two *in vitro* models^7,8^ by correlation analysis (shown in pink rectangle). BP: biological process, UMAP: Uniform Manifold Approximation and Projection.

Subsequently, we evaluated the DEGs in the oligodendrocyte and OPC clusters between NMS and MyS at weeks 8 and 15 (using an adjusted p-value < 0.05 and a log2 fold-change > 1) (Table S2). We observed stronger transcriptional differences between NMS and MyS at week 8 compared to week 15. At week 8, 33 genes were upregulated in NMS-derived oligodendrocytes (e.g., *ADAM23*, involved in juxtaparanodal organization^26^, and *GAB1*, a PDGF effector regulating OPC differentiation^27^), while only two were upregulated in MyS-derived oligodendrocytes (*TMSB4X*, which promotes cytoskeletal stability and myelin protein phosphorylation^28,29^, and *MT-ND3*, a mitochondrial gene linked to increased metabolic demand during maturation^30–32)^. In OPCs, 118 and 88 genes were upregulated in NMS and MyS respectively. NMS showed enrichment of early-stage transcription factors (*SOX4*, *SOX9*, and *PARP1)*, known to maintain progenitor identity or inhibit terminal differentiation^33–35^. By contrast, MyS-derived OPCs showed higher expression of genes promoting myelin membrane synthesis and maturation (*PLPP4*, *UGT8*, *LRRK2*, and *TMEFF2)*^36–39^. By week 15, transcriptional differences had diminished, with only three upregulated genes in NMS-derived oligodendrocytes and none in MyS. However, MyS-derived OPCs continued to show a maturation bias, with upregulation of genes important for lipid processing and cholesterol homeostasis (*LRP1B)*^40^. Together, these findings confirm that MyS cultures promote a more advanced oligodendrocyte maturation compared to NMS.

Next, we assessed the transcriptomic similarity of cellular populations, including the oligodendrocytes, developed in our MyS and NMS brain models with fetal and adult *in vivo* brain tissues as well as with *in vitro* brain models. To achieve this, we performed a comprehensive integrated analysis, incorporating references from two fetal^18,19^ and three adult ^20–22^ sn/scRNAseq brains datasets, as well as two *in vitro* hPSC-derived brain datasets^7,8^ (Figure 2d). This indicates a strong transcriptional similarity across the datasets, suggesting that the cellular populations in our models exhibit transcriptional profiles closely resembling those of adult human brain cells (Figure S3). Notably, MyS cultures contained a substantially higher proportion of oligodendrocytes (12.2% vs 1.45% week 8 and 7.52% vs 1.84% week 15) and astrocytes (14.03% vs 11.02% week 8 and 34.22% vs 21.00% week 15) compared to NMS. Next, we performed a Spearman correlation analysis. By merging the adult and *in vitro* datasets, we found that MyS-derived oligodendrocytes exhibited higher transcriptomic correlation with adult brain samples (0.66-0.67^20–22)^ compared to Marton *et al.* (0.54-0.59)^7^ and Fagiani *et al.* (0.42-0.50)^8^ *in vitro* datasets (Figure 2e). This finding was further supported by a comparative analysis of pseudobulk oligodendrocyte profiles visualized via scatter plots (Figure S4a-d).

We next performed Slingshot trajectory analysis on the integrated dataset to position oligodendrocytes derived from MyS and NMS within a developmental context. Using a supervised approach, we designated the youngest (9-12 weeks) fetal OPCs as the starting point^18^. This revealed three distinct lineages (Figure 3a). In the first lineage, MyS-derived oligodendrocytes appeared downstream of fetal OPCs and oligodendrocytes from the Marton *et al.* study^7^ but preceding mature adult oligodendrocytes^20–22^, suggesting a continuous trajectory toward maturation. The second and third trajectories did not extend to any adult oligodendrocyte populations. In the second trajectory, NMS and MyS oligodendrocytes were located very close together but downstream of oligodendrocytes from Marton *et al.*^7^, and fetal OPCs^18,19^. The third trajectory included all three *in vitro* created oligodendrocytes, with MyS-derived oligodendrocytes downstream of fetal OPCs. Collectively, these findings indicate that MyS-oligodendrocytes exhibit greater maturation compared to fetal OPCs and cells from other *in vitro* models.

**Figure 3.**
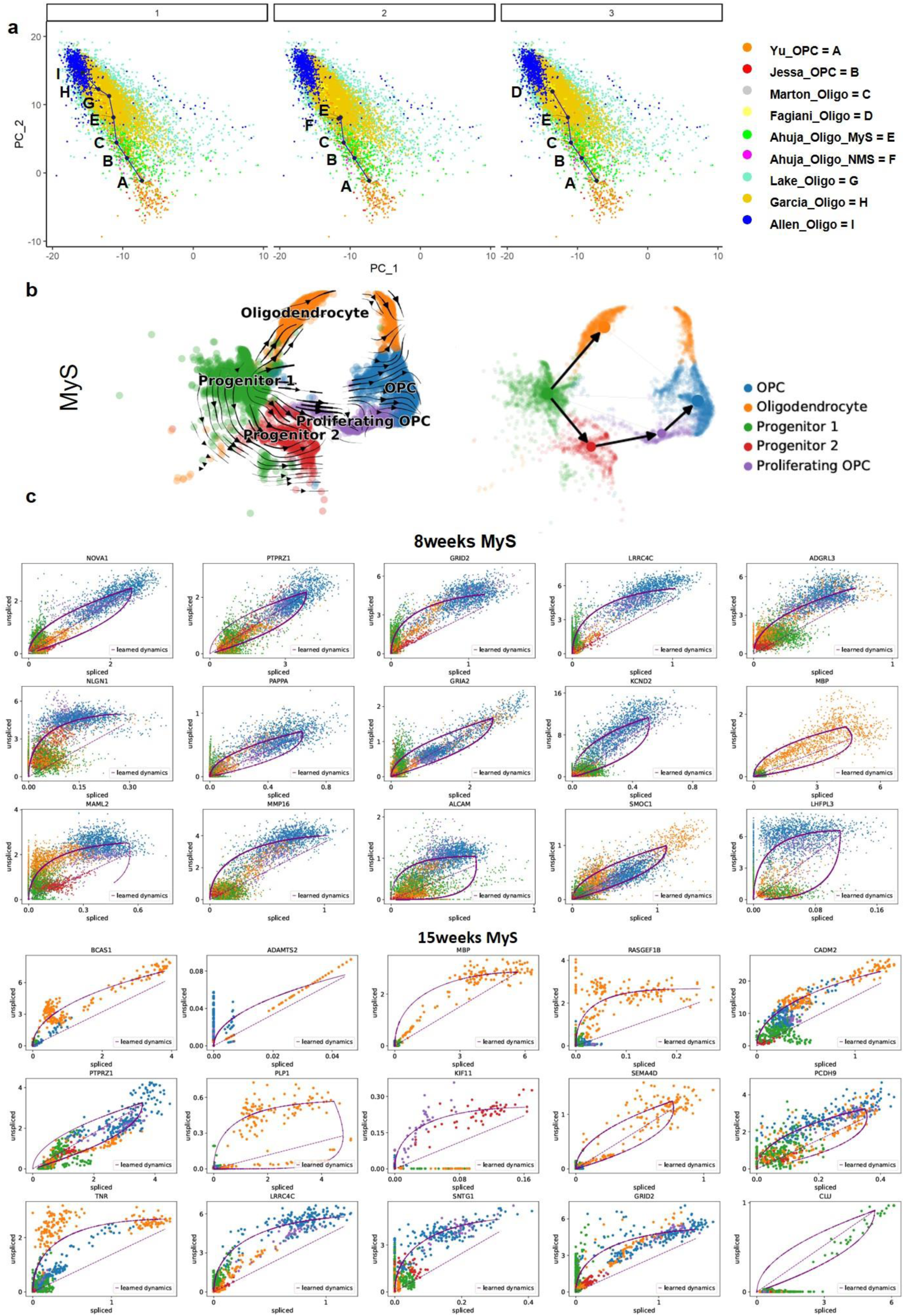
Enhanced maturation of MyS-derived oligodendrocytes compared to NMS, with significant progress observed at 15 weeks. **a** Slingshot analysis of integrated sn/scRNAseq datasets identified three distinct lineages. The first lineage included mature oligodendrocytes, with MyS-derived oligodendrocytes leading the developing OPCs from fetal datasets^18,19^ and oligodendrocytes from the *in vitro* model in Marton *et al.*’s^7^ study and followed by mature oligodendrocytes^20–22^. The second and third lineages consisted of developing OPCs and oligodendrocytes from *in vitro* brain models. Notably, in the second lineage, MyS-derived oligodendrocytes preceding NMS-derived oligodendrocytes. **b** Left: Velocity projection onto the UMAP embedding based on the dynamical model revealed two developmental trajectories starting from progenitor cells either directly to oligodendrocytes or via OPCs in MyS. Right: The PAGA velocity graph, overlaid on the UMAP embedding, provides a visualization of potential transitions between cell types. In this representation, nodes denote cell groups, while edge weights measure the connectivity between these groups. **c** Genes from MyS-derived cells at 8 and 15 weeks that exhibit significant dynamic behavior, positioning them as potential key drivers of the primary processes within certain cell populations. These genes were systematically identified through their characterization by high likelihoods in the dynamic model^25^.

To further explore the temporal dynamics of oligodendrocyte development, we performed RNA velocity analysis on week 8 and 15 MyS-derived progenitor, OPC, and oligodendrocyte clusters. The analysis conducted on pooled cells at 8 and 15 weeks MyS, revealed two developmental routes – one progressing directly from progenitors to oligodendrocytes, and another transitioning via OPCs (Figure 3b). Dynamic gene expression profiling revealed distinct gene sets at weeks 8 and 15, likely acting as key regulators of oligodendroglial maturation within defined cellular states^41^ (Figure 3c). Notably, clustering of OPCs and oligodendrocytes in week 15 MyS became more prominent along the developmental trajectory compared to those in week 8 MyS, indicating progressive maturation of the oligodendroglial lineage in MyS over time (Figure S5a–b, Table S3-4). At week 8, we found highly dynamic genes involved in early lineage commitment, synaptic signaling, and initial onset of myelin-related gene expression (*PTPRZ1*, *GRIA2*, and *KCND2)*^7,42,43^. Notably, *MBP* expression emerged at this stage, marking the onset of myelin gene activation. By week 15, several highly dynamic genes associated with MyS-derived oligodendrocytes were identified, exhibiting a transcriptional profile indicative of advanced maturation (*BCAS1*, *PLP1*, and *MBP*), which are crucial for active myelination and myelin compaction^44–46^. The continued presence of *PTPRZ1* in OPCs and *ADAMTS2* in oligodendrocytes further suggests roles in structural maintenance of mature oligodendrocytes^47^. These results support a temporal progression in MyS cultures from early OPCs toward myelinating oligodendrocytes.

### 3. Temporal development of myelin sheaths in MyS

We further analyzed oligodendrocyte maturation in MyS versus NMS by immunocytochemistry and transmission electron microscopy (TEM). Immunostaining of MyS demonstrated that neurons (TUBB3^+^), astrocytes (GFAP^+^), and elaborately branched oligodendrocytes (MBP+) were positioned in close proximity, with interwoven processes and complex morphologies suggestive of active cell-cell interactions (Figure 4a). This spatial arrangement was further supported by 3D rendering of the immunofluorescence images, which highlighted the intricate architecture and contact points between these cell types (Figure 4b). By contrast, the number of MBP⁺ oligodendrocytes in NMS was markedly lower, resulting in very few observable neuron-oligodendrocyte interaction sites (data not shown). TEM revealed that myelinated axons were already present in MyS at six weeks of culture (Figure 4c-d, Video S1 at week 10). To evaluate myelin maturation over time, we quantified multiple structural parameters on loose, mid-compact and compact structures in 10- and 15-week-old MyS, including G-ratio, axon radius, number and thickness of myelin lamellae per axon, and the percentage of different myelin types. In week 10 and 15 MyS, the G-ratio was lowest in loosely organized myelin structures (∼0.5), compared to mid-compact (∼0.72) and compact (∼0.66) myelin, indicating increased axonal wrapping with advancing myelin compaction (Figure 4e). Compact myelin sheaths containing more myelin lamellae with lesser thickness, were chiefly found around lower-diameter axons. By contrast, mid-compact myelin structures having fewer but thicker myelin lamellae were found around bigger axons, which was not different in week 10 or week 15 MyS (Figure 4f-g). The myelin lamellar thickness was the highest in loose myelinated axons compared to mid-compact and compact myelinated axons. However, myelin lamellar thickness was lower in week 15, compared to week 10 mid-compact myelinated axons, indicating myelin compaction over time in MyS (Figure 4h). Further, the percentage of mid-compact myelinated axons was higher than loose myelinated neurons in week 15 compared to week 10 MyS (Figure 4i). Consistent with the presence of a small number of MBP^+^ oligodendrocytes in NMS (Figure 1f), we observed limited myelinated axons in week 10 NMS (Figure S6a), supporting more robust and organized myelin formation in MyS.

**Figure 4:**
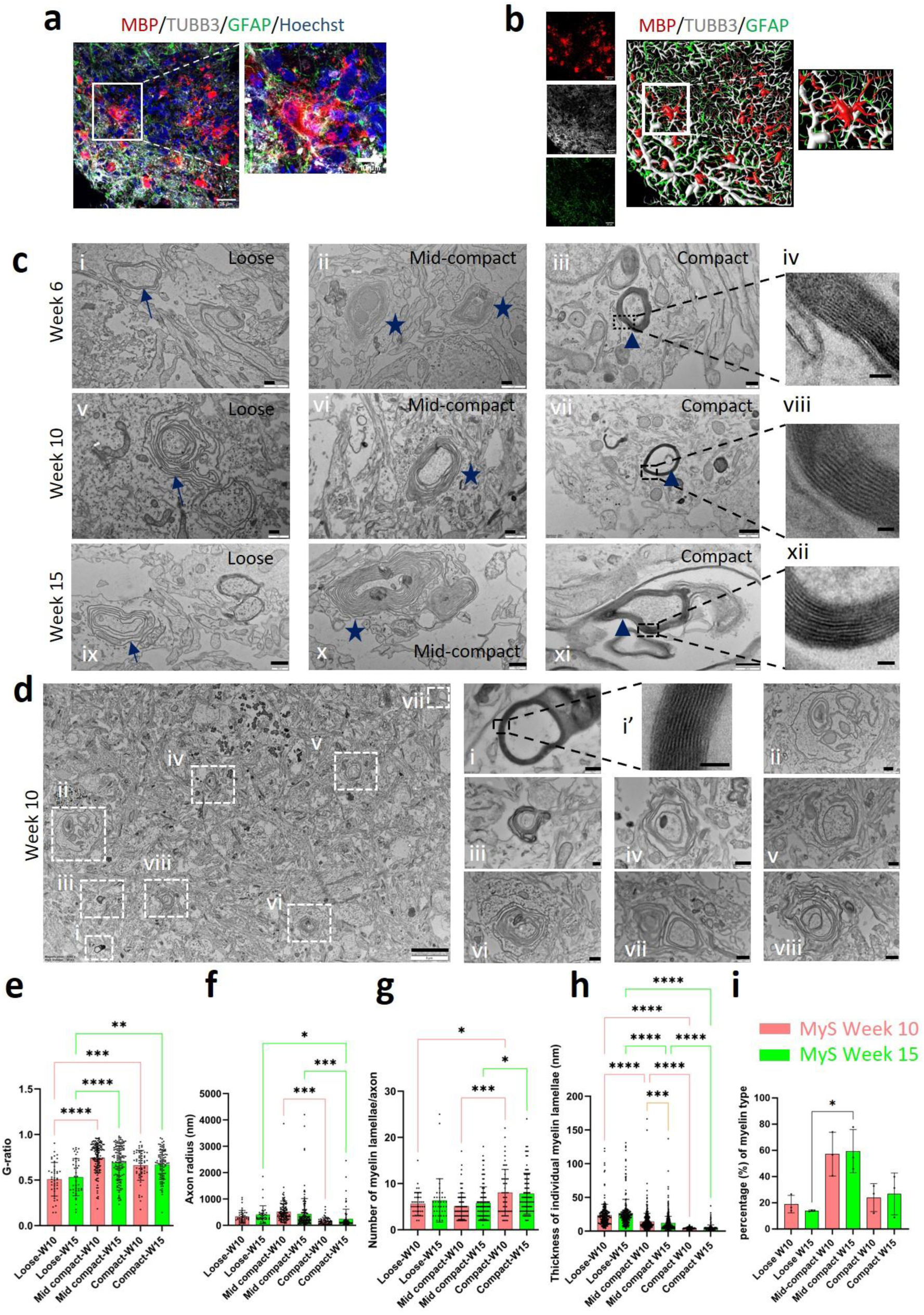
Oligodendrocyte maturation leading to formation of myelin sheaths in Sigma-iPSC0028 derived MyS. **a** Representative confocal image showing immunostained MBP^+^ oligodendrocyte processes interacting with TUBB3^+^ neurons and GFAP^+^ astrocytes in MyS at week 10. Nuclei were counterstained with Hoechst. **b** 3D rendering of confocal reconstructed image indicating TUBB3^+^ neurons surrounded by MBP^+^ oligodendrocyte and GFAP^+^ astrocytes processes in MyS at week 10. **c** Representative TEM images showing formation of different subtypes of myelin sheaths (loose (arrow) i, v, ix), mid-compact ((star) ii, vi, x) and compact ((triangle) iii, iv, vii, viii, xi, xii) in MyS at week 6,10 and 15 (scale bar = 500 nm: i, ii, v, vi, vii, ix, x; 200 nm: iii, xi; 45 nm: iv, xii; 25 nm: viii). **d** Representative TEM images showing distribution and types of myelin sheaths in MyS at week 10 (scale bar = 5 μm, 200 nm = i, iii; 45 nm = i’; 500 nm = ii, iv-viii). **e-i** Quantification of myelin sheath parameters: G-ratio, axon radius, number of myelin lamellae/axon, thickness of myelin lamellae and percentage of myelin type in MyS at week 10 and 15. Statistical analyses were performed by Two-way ANOVA with Sidak’s comparison test (**e-i**). Data shown are mean ± SD. (**a-d**) Representative TEM images from N=3 replicates, (**e-i**) N=3 (at least 200 axons (n>200) for all parameters) for myelin quantification. *p< 0.05, **p< 0.01, ***p< 0.001, ****p0.0001.

We also created NMS and MyS from human H9-ESC line derived NPCs and human i*SOX10*-H9-ESC OPCs to study stem cell line independent myelination. This revealed a similar pattern of oligodendrocyte presence using immunostaining for neurons (TUBB3^+^) and oligodendrocytes (MBP^+^) (Figure S6b-week 10), as well as myelination by TEM (Figure S6c-week 6, 10 and 13 and Video S2-week 10) in H9-ESC derived MyS. Again, only few myelinated axons were observed in H9-ESC derived NMS (data not shown).

### 4. Lipidomic profile supports myelin maturation over time in MyS

Lipids, especially cholesterol, are essential for CNS myelin sheaths, providing structural integrity, insulation, and facilitating rapid nerve impulse conduction. To demonstrate greater abundance of mature oligodendrocytes in MyS compared to NMS, a detailed lipidomic analysis was performed. The technical replicates of both NMS and MyS at weeks 8 and 15 clustered together in principal component analysis indicating technical reproducibility (Figure 5a), while heatmap visualization revealed substantial differences in lipid composition across the four groups (Figure S7a). We observed 17 lipid subclasses from which the relative amounts of phosphatidylcholine (PC) and phosphatidylethanolamine (PE) were the most represented in both NMS and MyS as reported in *in vivo* studies^48^ (Figure 5b).

**Figure 5:**
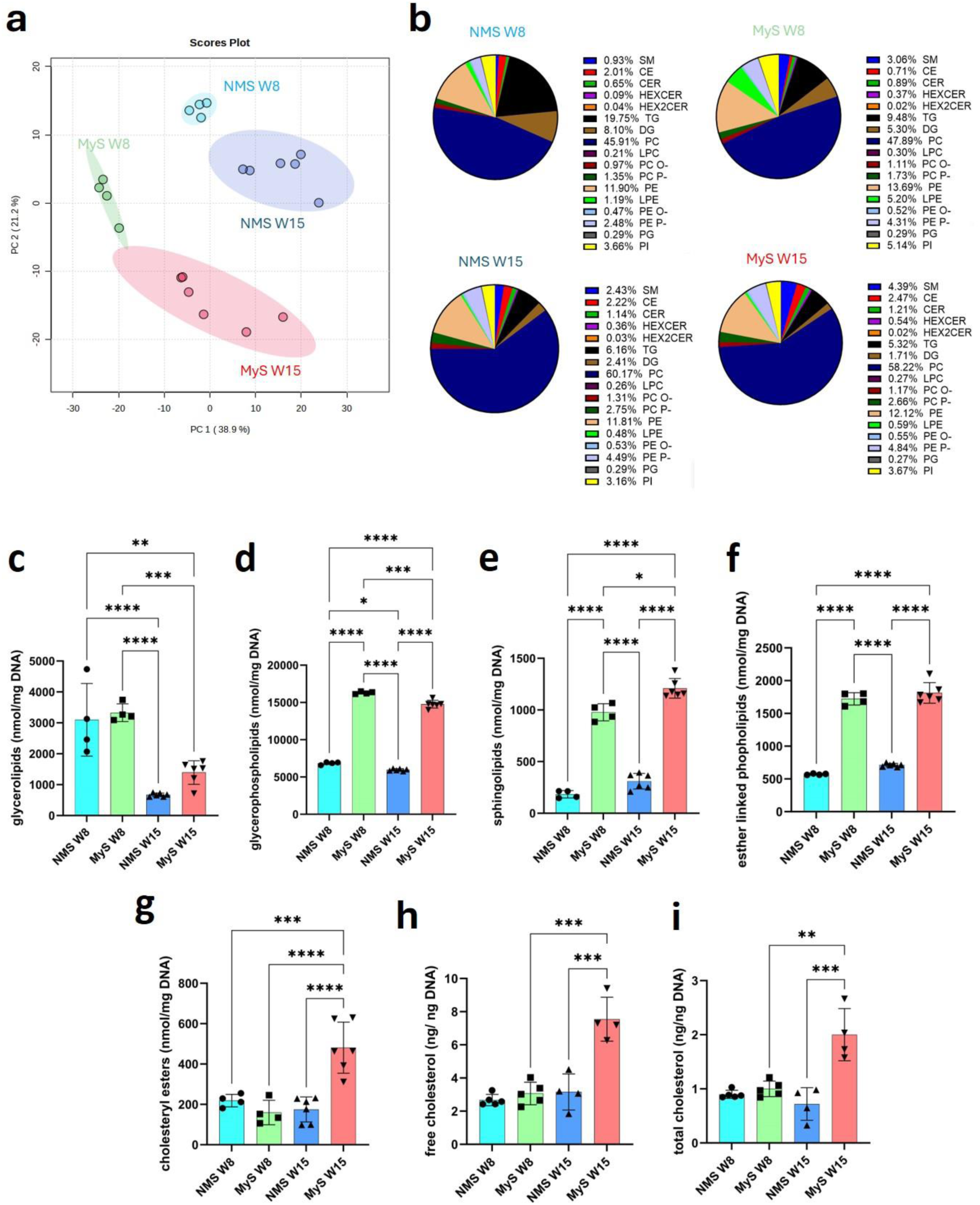
Temporal increase in myelin lipids in MyS suggesting maturation of myelin sheaths. **a** Pie charts indicating the different types of lipid species identified by lipidome analyses in NMS and MyS at week 8 and 15. **b** PCA analysis showing distinctive clustering of MyS and NMS samples at week 8 and 15. **c-g** Comparison of absolute amount of different major lipid classes identified in NMS and MyS at week 8 and 15 by lipidome analysis: glycerolipids, glycerophospholipids, sphingolipids, esther linked phospholipids, and cholesteryl esters. **h-i** Cholesterol (free and total) detection assay confirming the highest cholesterol amount in MyS at week 15 as compared to other conditions. Statistical analyses were performed by Two-way ANOVA with Sidak’s comparison test (**c-i**). Data shown are mean ± SD, n = 4-6. *p< 0.05, **p< 0.01, ***p< 0.001, ****p<0.0001. SM: Sphingomyelin, CE: Cholesteryl Esters, CER: Ceramides, HEXCER: Hexosylceramides, HEX2CER: Hexosyl-2-Ceramides, TG: Triglyceride, DG: Diglyceride, PC: Phosphatidylcholines, LPC: Lyso-Phosphatidylcholines, PC O-: Ether-linked Phosphatidylcholines, PC-P-: Plasmalogen Phosphatidylcholines, PE: Phosphatidylethanolamines, LPE: Lyso-Phosphatidylethanolamines, PE O-: Ether-linked Phosphatidylethanolamines, PE P-: Plasmalogen Phosphatidylethanolamines, PG: Phosphatidylglycerols, PI: Phosphatidylinositol.

To focus on changes in lipid fingerprint related to oligodendrocyte maturation and myelin sheath formation in NMS and MyS, we focused on five lipid classes: glycerolipids (triglycerides and diglycerides; Figure S7b-c), glycerophospholipids (e.g. phosphatidylcholine, lysophosphatidylcholine, phosphatidylethanolamine, lysophosphatidylethanolamine, phosphatidylglycerol, and phosphatidylinositol; Figure S7d-i), sphingolipids (including sphingomyelin, ceramides, hexosylceramides, hexosyl-2-ceramides; Figure S7j-m), esther linked phospholipids (Figure S7n-q), and cholesteryl esters. Glycerolipid levels declined over time in both models but remained moderately higher in MyS compared to NMs at both 8 and 15 weeks (Figure 5c). A similar trend was observed for glycerophospholipids, which showed a slight decline over time but remained higher in MyS compared to NMS at both time points (Figure 5d). These patterns suggest a possible metabolic shift toward higher myelin lipid synthesis and incorporation into myelin membranes in MyS compared to NMS. Sphingolipids remained stable in NMS but increased slightly in MyS over time, with overall levels higher in MyS across both weeks (Figure 5e). Esther linked phospholipids showed no significant temporal change in either model but were consistently higher in MyS than NMS (Figure 5f). Cholesteryl esters were notably higher in week 15 MyS compared to both week 8 MyS and NMS, indicating increased lipid storage or altered cholesterol metabolism at this stage (Figure 5g). Given the essential role of cholesterol in myelin sheath stabilization and compaction^49,50^, we further measured cholesterol levels using a fluorometric assay. Both free and total cholesterol were significantly elevated in week 15 MyS compared to all other groups, further supporting myelin maturation and compaction (Figure 5h– i). Together, these findings point to a more mature and metabolically active myelination profile by time in MyS compared to NMS.

### 5. MyS are functionally mature compared to NMS

To assess if myelinating oligodendrocytes influence neuronal activity, we employed high density-multielectrode array (HD-MEA) technology. Week 8-10 MyS and NMS were sectioned and cultured with the sliced surface positioned on HD-MEA plates (Figure S8a). In MyS, spontaneous spiking activity was detected as early as week 4 after plating the MyS slices. Based on the activity maps of MyS slices plated at 1-2 slices per well (Figure S8b), we applied a threshold of ≥150 electrodes with a firing rate >1 Hz to define activity, identifying 23 active and 10 non-active slices. Detailed analysis of neuronal activity from week 4 to 8 revealed unchanged spike amplitude, along with a significant increase in firing rate and number of active electrodes, as well as a shorter mean interspike interval, indicating enhanced network excitability and maturation (Figure S8c-f). While burst rate showed a non-significant upward trend, the accompanying significant increase in interburst interval further supports progressive network engagement over time (Figure S8g-h). To address variability in network behavior over time, we applied an additional cutoff of a ≥30% increase in overall spiking activity between weeks 4 and 8 to classify the slices by activity trend. Based on this threshold, number of active electrodes and firing rate of eight slices (34.8%) maintained activity levels (Figure S8i), three slice (13%) showed a decreasing trend (Figure S8j), 12 out of 23 slices (52.2%) exhibited an increasing trend from week 4 (Figure S8k), also evident in the raster plot at week 4 and 7 (Figure S8l). Conversely, no significant spiking and bursting activity was detected in the NM slice cultures over the 8 weeks of electrophysiology recording (data not shown).

### 6. MyS to model CNS disorders

We next assessed if the MyS model could be used to study central nervous system (CNS) diseases. As proof of concept for studying oligodendrocyte injury, we treated week 10 MyS with toxic phospholipid lysolecithin for 15 hours, which has been demonstrated to cause demyelination in *ex vivo* explant cultures^51^ and an *in vitro* myelination model^7^. Week 10 MyS treated with lysolecithin seemed to disintegrate as cells were shed from the spheres (Figure S9a). Immunostaining demonstrated that lysolecithin-treated samples contained less MBP^+^ oligodendrocytes than untreated control MyS (Figure S9b-c). Consistently, TEM demonstrated that lysolecithin-treated MyS contained broken myelin sheaths (Figure S9d).

We also investigated if the MyS model could be used to study CNS infections, specifically if the model supports Rabies virus (RABV) infection and replication. Week 10 MyS were infected with two different RABV strains, i.e. mCherry-SAD-B19 and CVS-11 (Figure 6a). RT-qPCR showed a temporal increase in RABV viral RNA in the supernatant of MyS infected with either mCherry-SAD-B19 or CVS-11, reflecting successful viral replication (Figure 6b). Immunofluorescence microscopy indicated clear viral infection for both virus strains, as shown by the expressed mCherry in mCherry-SAD-B19 infected cultures and by immunostaining for RABV Nucleoprotein in CVS-11 infected cultures. Virus replication was observed in all cultures independent of the virus inoculum (30, 100 or 300 TCID_50_ per spheroid) (Figure 6c-d). Quantification of the fluorescence signals revealed significant amounts of infected cells for cultures infected with the mCherry-SAD-B19 and CVS-11 strains (Figure 6e). We also performed TEM to better evaluate the pathological features of RABV infection^52,53^. We demonstrated the presence of bullet-shaped viral particles (Figure 6f), mitochondria with disintegrated cristae (Figure 6g)^54^, and degenerated cellular debris (Figure 6h). In addition, demyelinated neurons were observed, suggesting disruption of neuron-glia interactions either due to degeneration of neurons, glial cells or both (Figure 6i). We found disrupted neuronal morphology upon infection with CVS-11 but not with mCherry-SAD-B19, correlating with the higher pathogenicity of the CVS-11 RABV strain as compared with the attenuated SAD-B19 vaccine strain, as also shown in mice^55^ (Figure S11a). As MyS contain not only neurons, but also astrocytes and oligodendrocytes, we explored if the model would allow the study of RABV cellular tropism. The mCherry-expression for the mCherry-SAD-B19 strain was primarily found in neurons (NeuN^+^) and oligodendrocyte precursors (OLIG2^+^) but not in mature oligodendrocytes (MBP^+^) (Figure S11b). For the CVS-11 RABV strain, we observed colocalization of immunostaining for the viral N-protein with neurons (NeuN^+^), OPCs (OLIG2^+^) and oligodendrocytes (MBP^+^) (Figure S11c). Interestingly neither CVS-11 nor SAD-B19 appeared to colocalize with astrocyte markers (SOX9^+^/GFAP^+^). More extensive studies are required to quantify strain-dependent RABV infection.

**Figure 6:**
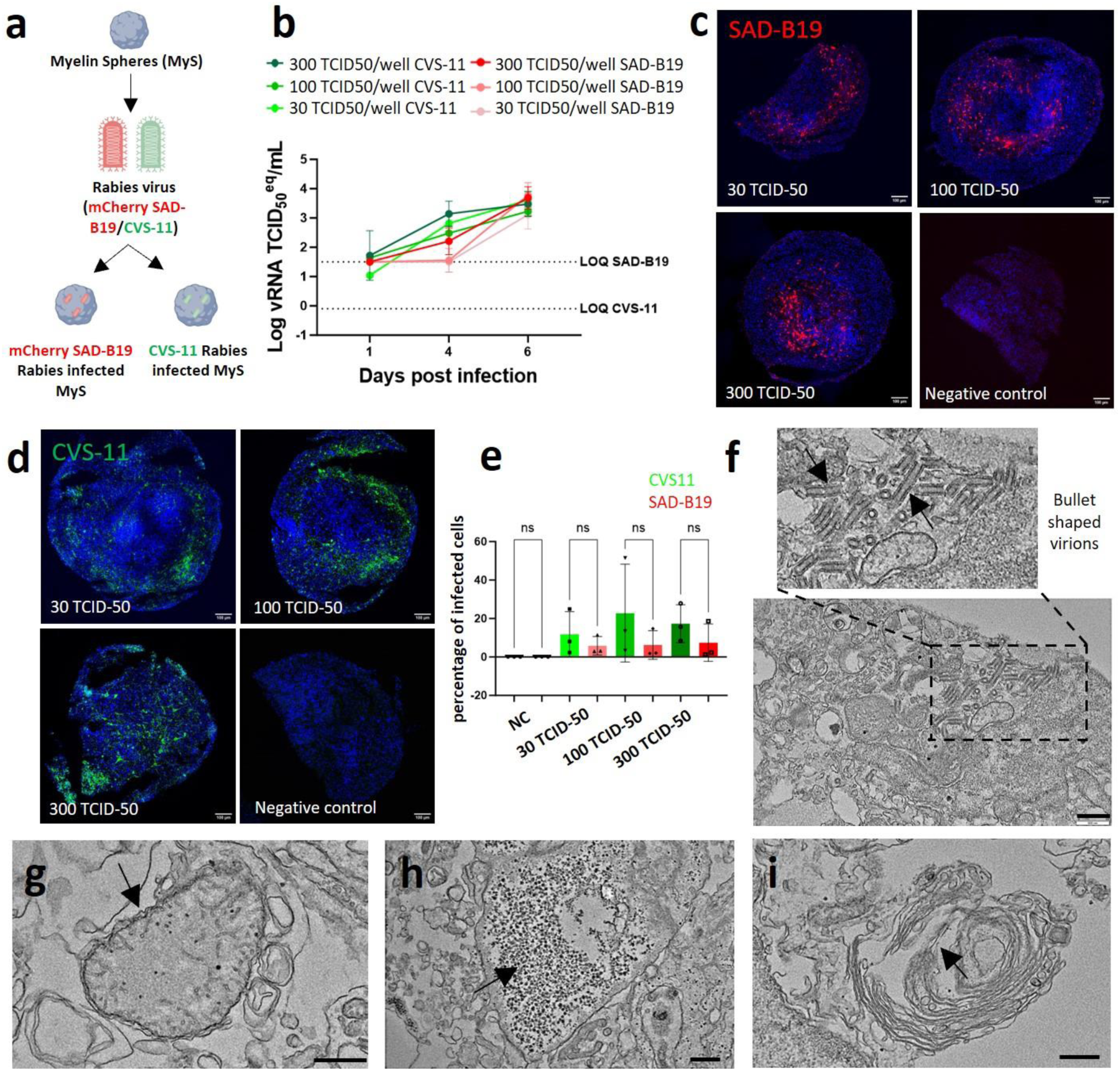
Rabies infection pathology was recapitulated *in vitro* using MyS. **a** Schematic representation of Rabies virus strains (mCherry-SAD-B19 and CVS-11) infection in week 10 MyS. **b** RT-qPCR showing increasing amount of viral RNA by days post infection for both mCherry-SADB19 and CVS11 infection in the supernatant of infected week 10 MyS. Viral RNA was quantified relative to a standard curve of known amounts of viral infectious units (TCID_50_ equivalents of viral RNA (TCID_50_^eq^)). LOQ = Limit of Quantification. **c-d** Fluorescent imaging of mCherry-SAD-B19 and immunostained CVS-11 at three viral concentrations (30, 100 and 300 TCID-50) in week 10 MyS at 5 days post infection (scale bar = 100 μm). **e** Quantification of fluorescence intensity of mCherry-SAD-B19 and CVS-11 in week 10 MyS at 5 days post infection with different viral concentrations. **f-i** Transmission electron micrographs showing several hallmarks of Rabies virus infection like bullet shaped virions (arrow, **f**), swollen mitochondria with disintegrated cristae (arrow, **g**), cell debris accumulation (arrow, **h**), axon demyelination (arrow, **i**). Scale bar = 500 nm. Statistical analyses were performed by Two-way ANOVA with Sidak’s comparison test (**e**). Data shown are mean ± SD. N=1, n=3-7 (**b**), representative images and quantification from N=3-4 (**c**, **d**, **f-i**), N=3 (n=3, each data point represents 3 slices **e**). ns = statistically not significant.

## Discussion

In this study, we developed hPSC-derived MyS by coculturing cortical NPCs with pre-differentiated i*SOX10*-OPCs to promote neuron-oligodendrocyte interactions and enhance *in vitro* myelination. Compared to NMS, MyS showed earlier and more robust myelination, with more oligodendrocytes appearing as early as six weeks. Over time, oligodendrocytes in MyS generated increasingly compact and multilamellar myelin sheaths and enhanced neuronal spiking activities, as confirmed by combined ultrastructural, lipidomic and electrophysiology evidence. Consistently, the transcriptomic profile of MyS-derived oligodendrocytes more closely resembles adult human brain-derived oligodendrocytes than those produced in other *in vitro* models^7,8^. Although MBP^+^ oligodendrocytes were also present in NMS, likely due to spontaneous differentiation in the presence of endogenous oligodendrocyte-specific growth factors as also reported in other studies^7,8^, their number was substantially lower. Notably, MyS also contained significantly more GFAP^+^ astrocytes and NeuN^+^ neurons compared to NMS. This suggests that inclusion of OPCs in MyS may influence the differentiation and maturation of other neural cell types, including astroglia and neurons. However, future studies will be required to further elucidate the mechanisms by which oligodendroglia might promote astroglial and neuronal differentiation, as well as to characterize the differences between MyS- and NM-derived astrocytes and neurons in greater detail.

SnRNAseq on MyS and NMS at week 8 and 15 confirmed the significantly greater percentage of oligodendrocytes (and astrocytes) in MyS. Evaluation of DEGs in OPCs and oligodendrocyte clusters at 8 and 15 weeks, as well as evaluation of highly transcribed genes from RNA velocity studies not only demonstrated much faster oligodendroglial cell development and maturation in MyS than NMS, but also a very clear temporal maturation pattern of MyS-derived oligodendroglial cells. We compared the maturation level of oligodendrocytes generated in MyS (and NMS) with those from other hPSC-based models^7,8^ and with oligodendrocytes from publicly available human brain datasets spanning fetal to adult stages^18–22^. MyS-derived oligodendrocytes showed a stronger correlation with adult brain oligodendrocytes across three snRNAseq datasets (correlation: 0.67-0.70), compared to those from the organoid-based model by Marton *et al.*^7^, which exhibited spontaneous myelination at DIV 100 (correlation: 0.58), and the Fagiani *et al.*⁸ model at DIV28-105, which used *SOX10* overexpression in NPCs^8^ (correlation: 0.47-0.49). The greater transcriptional maturity of MyS-derived oligodendrocytes, compared to those from NMS and other hPSC-derived models, is further supported by Slingshot analysis of the integrated sn/scRNA-Seq datasets. These analyses support the conclusion that oligodendrocytes generated in MyS cultures exhibit transcriptional profiles more closely resembling adult brain oligodendrocytes compared to those derived from other published hPSC-based models and the current NMS. Notably, the correlation indices between MyS-derived oligodendrocytes and oligodendrocytes generated in the other two PSC-derived *in vitro* models (>0.7) were relatively high, suggesting a shared *in vitro* phenotype. Further, MyS-derived oligodendrocytes also include cells resembling fetal OPCs, as seen in trajectories 2 and 3 – a pattern also observed in oligodendrocytes from Fagiani *et al.*^8^ and Marton *et al.*^7^. The reasons for the higher degree of transcriptional maturity of MyS-derived oligodendrocytes are likely multiple. While both the Fagiani *et al.*^8^ and MyS models employed *SOX10* overexpression to induce the oligodendrocyte lineage, key differences in study design likely contributed to the observed maturation differences. In Fagiani *et al.*, transcriptomic analysis was performed 8 weeks after *SOX10* transduction in NPCs, whereas MyS analysis occurred at week 8 or 15 after coculturing NPCs with 24-day-old i*SOX10*-OPCs – allowing for a longer maturation period. Furthermore, MyS spheroids incorporated >90% O4⁺ OPCs, as opposed to initiating *SOX10*-mediated commitment in NPCs. On the other hand, MyS and Marton *et al.*^7^ cultures had similar overall durations (80-129 vs. 100 days), yet only MyS included directed *SOX10* induction, underscoring the role of transcriptional programming and OPC pre-commitment in promoting more mature, adult-like oligodendrocyte development in MyS.

To further assess the development of mature oligodendrocytes, we performed detailed lipidomic analysis and cholesterol quantification in NMS and MyS cultures – an approach that, to our knowledge has not been previously applied to human *in vitro* myelination models. Consistent with the snRNAseq data, we detected significantly greater levels of lipids associated with myelin maturation and compaction in MyS. Notably, sphingolipids, essential for myelin sheath formation and stabilization^56^, increased over time in MyS, while glycerolipids and glycerophospholipids decreased, suggesting a metabolic shift toward specialized myelin lipid production. Furthermore, the observed increase in cholesteryl esters, as well as free and total cholesterol, in week 15 MyS relative to both week 8 MyS and week 8 or 15 NMS provides strong evidence for progressive myelin membrane compaction. These lipidomic findings are in line with the RNA velocity analysis, revealing greater activity of genes involved in myelin compaction and terminal oligodendrocyte maturation at week 15.

Presence and maturation of myelin in MyS was corroborated by TEM analysis. The G-ratio of myelinated axons in MyS (0.5-0.75) was in the range seen in the adult human cortex (0.5-0.6), suggesting that myelination in MyS recapitulates to a certain extent the *in vivo* developmental process^57,58^. Nevertheless, the number of myelin lamellae per axon (i.e. 5-8 lamellae) in MyS was still more fetal-like, as myelinated axons in adult human brain cortex typically contain 10-50 myelin lamellae^59^.

The measured neuronal activities in MyS, but not in NMS, by HD-MEAs suggest the functionality of the myelinating oligodendrocytes in MyS. However, as we did not perform extensive immunostaining controls for recorded NM slices, we cannot exclude that neurons from NMS are more fragile and may be more prone to damage during medium changes. It should also be noted that only ±75% of MyS slices were functionally active, and that an increase in temporal electrophysiological activity was only seen in ±50% of these slices. This variability may stem from biological differences in neuron-glia interactions across spheroids or from technical limitations, such as inconsistent attachment of MyS slices to HD-MEAs. In the future, this could be assessed by immunostainings for neurons, astrocytes and oligodendrocytes; however this approach is difficult in longitudinal studies unless lineage-specific fluorochromes are introduce in hPSCs. In addition, the random 3D alignment of axons in MyS makes it nearly impossible to evaluate features such as the effect of myelination on conduction velocity. This problem could be resolved by culturing MyS in scaffolds structures or microfluidics, which would align the axons in one direction. Combined with HD-MEAs, this would enable stimulating soma and detecting the electrical signals at the level of axons.

Lastly, to explore the applicability of the MyS model for studying neurological diseases, we established two pathological paradigms – (1) lysolecithin-induced demyelination and (2) RABV infection. Marton *et al.*^7^ demonstrated that lysolecithin decreases the percentage of SOX10 positive oligodendroglia in their hPSC-derived cultures, in line with our data on structural deformation of myelin sheaths using TEM. Rabies is a fatal neurotropic disease with a >99.9% mortality rate once neurological symptoms appear. We demonstrated successful RABV infection of MyS via immunofluorescence and TEM, and observed some degree of strain-dependent neurotropism^60^. Notably, infection of astrocytes by the CVS-11 strain has been shown to depend on the route of administration in mouse models, underscoring how viral tropism is shaped by experimental conditions^61^. Furthermore, reports of RABV localization in Schwann cells^62^ suggest a capacity to target myelinating cells, although earlier studies offer conflicting evidence on this phenomenon^63,64^. While precise quantification of cell type-specific infection in the MyS model was difficult, due to tight cell-cell contacts, immunostaining did not demonstrate infection of astrocytes by the pathogenic CVS-11 or the attenuated SAD-B19 strain. In addition, MBP^+^ oligodendrocytes appeared to only be infected by CVS-11, while OLIG2^+^ OPCs appeared to be infected by both viral strains. These findings highlight the relevance of the MyS model for investigating glia-virus interactions in a controlled, human-derived *in vitro* setting. Future studies should integrate sn/scRNAseq for better neurotropism studies.

There are some limitations to the MyS model that should be acknowledged. Unlike conventional brain organoids, MyS do not recapitulate the typical layered cortical architecture of the human cortex. Although the transcriptome of MyS-derived oligodendrocytes resembles adult transcriptional profiles more closely than oligodendrocytes described in other *in vitro* models, incomplete maturation remains a concern. In line with this, ultrastructural studies suggest a late fetal state of myelination, not yet an adult brain oligodendrocyte myelination phenotype. Obviously, the transcriptomic differences between MyS and *in vivo* oligodendrocytes may be in part caused by *in vitro* culture artifacts, which could be better understood by grafting iPSC-derived OPCs into *in vivo* models, as has been done for other cell types (e.g., microglia^65^). Although there is some evidence that presence of maturing oligodendrocytes may allow electrophysiological maturation, the incompletely understood variability between different spheroid slices remains an obstacle, and additional testing models may be needed to corroborate these findings. Finally, while our data suggest enhanced astrocyte generation in MyS, this requires more in-depth validation of the snRNA-seq analysis.

Despite these limitations, the MyS model is a major advancement over current *in vitro* myelination models as a relatively fast and robust human-relevant myelination platform. It holds potential for the evaluation of compounds that promote oligodendrocyte differentiation and myelin repair. The cellular diversity present also provides opportunities to study cell-specific tropism in a tractable human model, with potential for extending this beyond RABV to other brain-tropic viruses; as well as anti-viral drug screens. Additionally, the MyS system is modular in design, allowing for disease modeling by mixing healthy and mutant NPCs or OPCs. This should make it possible to dissect cell-autonomous vs. non-cell-autonomous effects in de-myelinating disorders such as MS, and in other neurodegenerative disorders where defects in oligodendrocytes at least contribute to neurodegeneration^66^.

## Methodology

### Culture of hPSCs

Two hPSC lines, Sigma-iPSC0028 (iPSC EPITHELIAL-1, Sigma, female, RRID: CVCL_EE38) and H9 embryonic stem cells (H9-ESCs) (WA09 WiCell research Institute, female, RRID: CVCL_9773) were used in this study. The hPSCs were genome engineered with an inducible transcription factor *SOX10* cassette in the Adeno-associated virus integration site 1 *(AAVS1)* locus via zinc finger nuclease gene editing to drive effective differentiation towards oligodendrocytes. hPSCs, with or without inducible *SOX10* (i*SOX10*) cassette were maintained on hESC qualified Matrigel (Corning) coated 6-well plates in E8 flex medium (E8 basal medium with E8 supplement flex and 1% penicillin/streptomycin (Gibco)). hPSCs were passaged twice a week with 0.5 mM ethylenediaminetetraacetic acid (EDTA). The cells were regularly tested for mycoplasma contamination using the MycoAlert Mycoplasma Detection Kit (Lonza).

### Differentiation of hPSCs to cortical NPCs

Cortical NPCs were generated following the protocol described by Shi *et al.*^9^. Sigma-iPSC0028 and H9-ESCs were cultured on Matrigel-coated 6-well plates (Corning) in mTESR (StemCell Technologies) with Revitacell (Life Technologies). hPSCs were allowed to proliferate to 90% confluency, dissociated into single cells by Accutase (Sigma) and seeded at 2.5 million cells per well of a 6-well plate in neural induction medium (NIM, Table S5). The medium was changed every day for 11 days. On day 12, the neuroepithelial cells were dissociated with Dispase II (Sigma), cultured for an additional 4 days with neuronal maintenance medium (NMM, Table S5) with 20 ng/mL basic fibroblast growth factor (bFGF) and purified by passaging rosette-forming neuroepithelial cells twice with Dispase II (one each after 5-7 days). NPCs were dissociated on day 33 with Accutase and cryopreserved in NMM complemented with 10% dimethyl sulfoxide (DMSO) (Sigma) for further experimental use.

### Differentiation of hPSCs to OPCs

OPCs were generated by overexpression of the transcription factor *SOX10* in Sigma-iPSC0028 and H9-ESCs cell lines as described in Garcia *et al.*^10^. Briefly, Sigma-i*SOX10*-iPSC0028 iPSCs and i*SOX10*-H9-ESCs were cultured in E8 flex medium, dissociated with Accutase and plated at a density of 25,000 cells/cm^2^ on a Matrigel-coated 6-well plate. After 1 day, oligodendrocyte induction medium (OIM, Table S5) was added for the following 6 days. On day 7, OIM containing 1 μM Sonic hedgehog agonist (SAG) was added and the medium was refreshed daily until day 11. Cells were then dissociated with Accutase and replated at 50,000-75,000 cells/cm^2^ in the Basal Medium (Table S5) on poly-L-ornithin (PLO)-laminin coated 12-well plate to direct them towards oligodendrocyte fate. The medium was changed every day until day 23. On day 24, the cells were dissociated with Accutase and cryopreserved in the same medium with 15% DMSO for further experimental use.

### Generation of NMS and MyS

Days *in vitro* (DIV) 33 cortical NPCs were thawed in NMM supplemented with Revitacell (Life Technologies) and maintained in culture for five days. On the day of sphere generation, DIV24 OPCs were thawed in NMM and DIV38 NPCs were dissociated with Accutase followed by cell counting by nucleocounter (ChemoMetec). 3D spheres were generated by diluting the cells in 2X NMM (NMM supplemented with 0.5X B27 (Gibco), 0.5X N2 (Gibco)), 20% KnockOut Serum Replacement and 1% methylcellulose (Sigma) and forming 30 μL droplets hanging onto the inside part of a turned-over lid of a petri dish. The 3D NMS and 3D MyS comprised of 90,000 DIV38 NPCs or 30,000 DIV38 NPCs with 60,000 DIV24 OPCs, respectively, per droplet. The hanging drops were carefully transferred to the incubator for 24 hours, at which time the aggregated spheres were collected with a cut P1000 tip and transferred to 12-well ultra-low attachment plates (Corning) containing oligodendrocyte maturation medium (OMM, Table S5) supplemented with 3 μg/mL doxycycline for a week followed by only OMM. Plates were maintained on a rotating platform (IKA Wobble shakers Rocker 3D) on 50 revolutions per minute (rpm) for 6-15 weeks.

### Quantification of gene expression by real-time quantitative polymerase chain reaction (RT-qPCR)

DIV 33 NPCs and DIV 24 OPCs monocultures were collected as a cell pellet respectively, and RNA extraction was performed by Quick-RNA Microprep Kit (Zymo research). The total amount of RNA was quantified using the NanoDrop system (Thermo Fisher Scientific) and 500 ng to1 μg RNA was used to prepare cDNA using SuperScript^TM^ III First-Strand Synthesis System kit (Thermo Fisher Scientific). For 3D cultures, two spheroids of MyS or NMS were combined for each sample. RNA extraction, quantification and cDNA preparation were performed as mentioned above. RT-qPCR was performed to assess the expression of neuronal, astroglial and oligodendrocyte lineage markers by using Platinum SYBR Green qPCR Supermix-UDG (Thermo Fisher Scientific) on the ViiA 7 Real-Time PCR System (Applied Biosystems, USA). Analysis of gene expression was done by normalizing the cycle threshold (CT) value to that of the housekeeping gene Glyceraldehyde-3-phosphate dehydrogenase (GAPDH) and the delta CT was plotted. Primer sequences for each marker are listed in Table S6.

For Rabies virus (RABV) infected 3D MyS, RNA extracted from the culture supernatant (MACHEREY-NAGEL, cat 740956) was analyzed by RT-qPCR with iTaq™ Universal SYBR® Green One-Step Kit (BIO-RAD). Primers specific for RABV SAD-B19 and CVS-11 are listed in Table S6. The standard curve was generated from the RNA extracted from a 1/10 serial dilution series of a virus stock with known amount of infectious units of 3.6 × 10^7^ TCID_50_/mL (determined using end-point dilution and expressed as tissue culture infected dose 50% (TCID_50_)). A 20 μL qPCR reaction comprised 4 μL of extracted RNA or standard, 10 μL of iTaq Universal SYBR® Green reaction mix, 0.25 μL of reverse transcriptase and 600 nM of each forward and reverse primer. RT-qPCR was performed on a QuantStudio 5 (Thermo Fisher Scientific) with the following protocol: 10 minutes (min) at 50 °C for reverse transcription, 1 min at 95 °C for polymerase activation and DNA denaturation, 40 cycles of 95 °C 15 seconds (s) and 62 °C 30s for PCR amplification. Viral RNA copies were quantified based on the standard curve and are represented as TCID_50_ equivalents/mL (TCID_50_^eq^/mL). The limit of quantification (LOQ) of viral RNA is defined by the lowest tested concentration from the standard curve that is still in the linear range.

### Immunofluorescence microscopy and quantification of cellular composition of NMS and MyS

Week 10 MyS or NMS harvested at different timepoints were fixed in 4% paraformaldehyde (PFA) overnight at 4 °C. The next day, PFA was replaced with 30% sucrose followed by overnight incubation at 4 °C. Spheroids were collected using a cut P1000 tip and embedded in Optimal Cutting Temperature (OCT) compound by snap-freezing in liquid nitrogen. Cryosectioning was performed using a CryoStar NX70 (Thermo Fisher Scientific) to obtain 30 µm-thick sections, which were stored at -20 °C until use. For immunostaining, 3-6 sections per spheroid (one central and two peripheral) were selected. Sections were washed with phosphate-buffered saline (PBS), blocked and permeabilized using 0.5% Triton X-100 in PBS (PBST) supplemented with 5% normal goat serum (Dako) for 1 hour at room temperature (RT). Samples were incubated with primary antibodies (Table S7) diluted in PBST overnight at 4 °C in humidified chambers, washed with PBS three times with 5 min interval, and secondary antibodies (species matched) diluted in Dako Real Antibody diluent were added for 1 hour at RT. Samples were washed with 0.01% PBST, three times with 10 min interval, and nuclei were stained with Hoechst 33342 (Sigma, 1:1000 dilution). Quantification of neurons, astrocytes, and oligodendrocytes was performed using QuPath software^11^. Cell types were identified based on marker-specific fluorescence intensity, and automated detection pipelines were used to segment and quantify positive cells within defined regions of interest (ROIs). Thresholds were optimized and kept constant to ensure comparability across samples.

RABV CVS-11 was kindly provided by Prof. Steven Van Gucht (Sciensano, Brussels, Belgium). RABV mCherry-SAD-B19 was kindly provided by the laboratory of Prof. Ashley C Banyard (Animal & Plant Health Agency, Surrey, UK). The mCherry sequence is inserted before the first gene within the genome of the SAD-B19 strain as described^12^. 3D MyS samples infected with RABV (CVS-11 or mCherry-SAD-B19) were sliced (30 µm thickness) using cryostat (Cryostar NX70, Thermo Fisher Scientific). The slices were incubated with an anti-RABV Nucleoprotein antibody (FITC Anti-Rabies Monoclonal Globulin (Table S7) diluted in PBST after nuclei staining for overnight at 4 °C, followed by thrice PBS washing. mCherry fluorescence was used as an indication for RABV infection in SAD-B19 (mCherry tagged) RABV infected 3D MyS samples (validated with the anti-RABV Nucleoprotein antibody in Figure S10). 3D MyS slices immunostained with cell type specific markers and/or RABV markers were mounted with ProLong Gold antifade mounting reagent (Life technologies). Confocal imaging was done using a Nikon C2 confocal microscope equipped with long-distance objective lens. 3D rendering images were made using the confocal images, afterwards using the VAA3D software (VAA3D-Neuron2_Autotracing)^13^.

### Transmission electron microscopy (TEM)

Spheroids were fixed in 2.5% glutaraldehyde (Electron Microscopy Services #16220) in 0.1 M sodium cacodylate buffer (Electron Microscopy Services #12300), pH 7.6 for 45 min on a shaker at 50 rpm at RT. Samples were washed thrice with 0.1M sodium cacodylate buffer and 1% osmium tetra-oxide (Electron Microscopy Services #19151) with 1.5% potassium ferrocyanide (Sigma #455989) diluted in 0.1 M sodium cacodylate buffer was added. Samples were incubated for an hour at 4 °C on a shaker. The samples were washed with Milli-Q water six times at 2 min intervals, followed by overnight incubation in 2% Uranyl acetate (Electron Microscopy Services #22400) diluted in Milli-Q water at 4°C. The next day, samples were washed five times for 7 min with Milli-Q water, before lead aspartate staining (Lead Nitrate-Electron Microscopy Services #17900 and L-aspartic acid-Sigma #11189, pH 5.5) for 30 min at 60 °C. Next, samples were washed with Milli-Q water thrice and dehydration was done by increasing concentrations of ethanol (30%, 50%, 70%, 90%, 100%) on ice, 10 min each. A second 100% ethanol step was done, followed by twice propylene oxide (Thermo Fisher Scientific #10487920) treatment for 10 min at RT. Spheroids were transferred to epon resin/propylene oxide mixtures before overnight infiltration with pure epon resin (Agar Scientific #AGR1081). The next day, the spheroids were embedded in inverted beam capsules and 70 nm thick slices were sectioned from the spheroids using the Leica Reichert Ultracut S2 and imaged using a JEM1400 transmission electron microscope (JEOL), operating at 80kV and equipped with an Olympus SIS Quemesa 11MP camera.

Quantification of myelin sheets in a 70 nm sections, one from the periphery (around 50 nm from the surface) and one from the center (around 200 μm from the surface) of each spheroid was combined and considered as one biological replicate. More than 220 axons from both week 10 and week 15 MyS samples were manually quantified across three independent biological replicates to assess various myelin sheath parameters. Measurements including axon radius, myelin thickness, and the number of myelin lamellae were performed using ImageJ on TEM images. Myelinated axons were classified into three categories based on interlamellar spacing: loose (>40 nm), mid-compact (15-40 nm), and compact (<15 nm) myelin. The G-ratio was calculated as the ratio of the inner axonal radius to the total outer radius (axon + myelin). Percentage for each myelin type was determined by dividing the number of axons in that category (loose, mid-compact or compact) by the total number of myelinated structures per biological replicate. Additionally, sections of 300 nm were cut with the Leica Reichert Ultracut S2 and tomograms were acquired using a Jeol 200kV S/TEM JEM F200 electron microscope with cold Field Emission Gun. Tilt series were obtained by SerialEM with a step width of 3° from -60° to 60° in STEM mode using a JEOL Bright Field detector. The 3D reconstruction was done with IMOD and the 3D representation with Amira software (Thermo Fisher Scientific, Video S1-2).

### Lipidomics

NMS and MyS Spheroids were collected at weeks 8 and 15. The spheroids were homogenized in 800 μL water with a handheld sonicator, and an amount of homogenate containing 1 μg DNA was diluted with water to 700 μL and mixed with 800 μL 1 N HCl:CH_3_OH 1:8 (v/v), 900 μL CHCl_3_ and 200 μg/mL 2,6-di-tert-butyl-4-methylphenol (BHT; Sigma-Aldrich), 3 μL of SPLASH® LIPIDOMIX® Mass Spec Standard (Avanti Polar Lipids, 330707), and 3 μL of Ceramides and 3 μL of Hexosylceramides Internal Standards (cat. no. 5040167 and 5040398, AB SCIEX). The organic phase was rated and evaporated by Savant Speedvac spd111v (Thermo Fisher Scientific) at RT for 1-2 hours. The remaining lipid pellets were stored at -80 °C for further use. For the DNA quantitation, a standard curve of herring sperm DNA concentrations in homogenization buffer (0.05 M NaHPO_4_/NaHPO; buffer, 2.0 M NaCl, 2.10 M EDTA, pH 7.4) was made. 10 μL of the sample homogenates was mixed with homogenization buffer and incubated for an hour at 37 °C in dark. 1 μg/mL Hoechst dye (Calbiochem) was added to both standards and samples, and fluorescence was measured at excitation filter of 360-390 nm and an emission filter of 450-470 nm for determination of DNA concentrations. Samples were assessed on a liquid chromatography electrospray ionization tandem mass spectrometer (Nexera X2 UHPLC system (Shimadzu) coupled with 6500+ QTRAP system; AB SCIEX) to identify several lipid classes. All lipid species were measured with a scheduled Multiple Reaction Monitoring (MRM) method. Sphingomyelin, cholesterol esters, ceramides, hexose-ceramides, and lactose-ceramides were measured in positive ion mode with the MRM transitions being based on fragment losses m/z 184.1, 369.4, 264.4, 264.4, and 264.4, respectively. Triglycerides, diglycerides and monoglycerides were measured in positive ion mode with the transitions being based on the neutral loss of one of the fatty acyl moieties. Phosphatidylcholine, alkylphosphatidylcholine, alkenylphosphatidylcholine, lysophosphatidylcholine, phosphatidylethanolamine, alkylphosphatidylethanolamine, alkenylphosphatidylethanolamine, lyso phosphatidylethanolamine, phosphatidylglycerol, phosphatidylinositol, and phosphatidylserine were measured in negative ion mode with MRM transitions for the fatty acyl fragment ions.

Peak integration was performed with the MultiQuant^TM^ software version 3.0.3. Lipid species signals were corrected for isotopic contributions (calculated with Python Molmass 2019.1.1). The total lipid amount and concentration per lipid class were expressed as absolute values in nmol/mg DNA. The lipidomic dataset was analyzed using the web-based application MetaboAnalyst 6.0^14,15^. Missing values were addressed through imputation. Lipid species with >50% missing values were removed, and the remaining missing values were substituted with the lowest of detection imputation method (1/5 of the smallest positive value of each variable). Data were normalized with a log_10_ transformation, and a Pareto scaling was applied (mean-centered and divided by the square root of the standard deviation of each variable). A further step was performed using the EigenMS method to estimate batch effects and identify systematic biases^16^. This normalization method employs a combination of analysis of variance (ANOVA) and singular value decomposition applied to the residuals matrix to remove these biases. Groups were compared using principal component analysis and hierarchical clustering heatmaps (distance measure: Euclidean, clustering algorithm: ward, standardization: auto-scale features).

### Cholesterol detection

Week 8 and 15 spheroids were collected, and lipids extracted by homogenizing the spheroids in 100 μL water with a handheld sonicator. DNA measurement was done by preparation of a standard curve of DNA measurements were done using Hoechst dye as mentioned above. Following DNA measurements, the samples were dehydrated using a speed-vacuum for an hour. The dry pellet was used for cholesterol measurement using the Cholesterol/ Cholesteryl Ester Assay Kit (Abcam, #ab65359) according to the manufacturer’s protocol. A total of 120 ng of DNA from each sample was used to measure cholesterol concentrations, and fluorescence was recorded using a 520 nm excitation and 580-640 nm emission filter.

### Sample preparation for single-nuclei RNA sequencing (snRNAseq)

For nuclei isolation, all steps were done on ice with pre-cooled buffers, and centrifugation was maintained at 4 °C. Briefly, week 8 or 15 old NMS and MyS (4-6 spheroids per sample) were transferred to 250 µL of ice-cold homogenization buffer (HB; 10 mM Tris pH 7.8, 0.1 mM EDTA, 320 mM sucrose, 5 mM CaCl_2_, 3 mM magnesium acetate, 0.1% NP40, 1X complete protease inhibitor (Roche), 1 mM β-mercaptoethanol, and RNase inhibitor (Promega)) for 3 min to preserve nuclear integrity. Samples were gently homogenized using pestle A (5 strokes) followed by pestle B (15 strokes) and the homogenates were passed through 70 µm EASYstrainer (Greiner). The homogenizer and the strainer were washed with additional 270 µL of HB, and 520 µL of gradient medium (10 mM Tris pH 7.8, 5 mM CaCl_2_, 50% Optiprep, 3 mM magnesium acetate, 1X complete protease inhibitor, 1 mM β-mercaptoethanol, and RNase inhibitor) was added to the samples. This solution was carefully layered onto a cushion solution (29% Optiprep, 150 mM KCl, 30 mM MgCl_2_, 60 mM Tris pH 7.8, and 250 mM sucrose) and centrifuged at 9000 rpm for 25 min at 4 °C. The supernatant was discarded carefully, and nuclei pellets were then resuspended in 25 µL of resuspension buffer (1X PBS, 2% bovine serum albumin (BSA)), filtered through Flowmi 40 µm cell strainers, and counted using a LUNA automated cell counter (Logos Biosystems) following the manufacturer’s instructions. The diluted nuclei suspensions were subsequently loaded onto the 10X Chromium Single Cell Platform (10X Genomics) (Next GEM Single Cell 3′ library and Gel Bead Kit v3.1) in accordance with the manufacturer’s protocol (10X User Guide; CG000315, Revision E). Steps including generation of gel beads in emulsion (GEMs), barcoding, GEM-RT cleanup, complementary DNA amplification, and library construction were all performed according to protocol. The quality of individual samples was checked with a Fragment Analyzer (Agilent), and library quantification was done using Qubit 4.0 (Thermo Fisher Scientific) before pooling the libraries. The final pooled library was sequenced on a NovaSeq6000 (Illumina) instrument at the VIB Nucleomics Core, using the Illumina NovaSeq 6000 v1.5 sequencing kit S4 200 paired-end reads (28-10-10-90), 1% PhiX with 100-base-pair paired-end reads.

### Quality control, snRNAseq data pre-processing

The Cell Ranger pipeline (version 7.1.0, 10x Genomics) was employed for sample demultiplexing and the generation of FASTQ files for read 1, read 2, and the i7 sample index of the gene expression library. Read 2 sequences from the gene expression libraries were aligned to the reference genome (Human GRCh38 (GENCODE v32/Ensembl98)) using the STAR aligner through CellRanger. Subsequent processing steps, including barcode processing, filtering of unique molecular identifiers (UMIs), and gene counting, were conducted using the Cell Ranger suite version 7.1.0.

### Normalization, clustering, and differential gene expression analysis of snRNAseq data

The gene count matrix for each sample was processed to exclude nuclei with fewer than 200 expressed genes and genes expressed in fewer than 3 nuclei. Additionally, quality control was performed using the Scater package (Version 1.32.1), both univariately and multivariately. This involved filtering out nuclei with extremely low or high total counts/transcripts (library size) (median absolute deviations (MAD) of 5), nuclei with low or high numbers of expressed genes (MAD of 5), and nuclei with a very high percentage of mitochondrial transcripts (MAD of 15). We used the Seurat pipeline (Seurat package, version 5.0.3) for normalization, scaling, and clustering. A Seurat object was constructed for each individual sample from the counts and metadata, followed by log-normalization of the count matrix. The most highly variable genes (HVGs) were identified using the FindVariableFeatures function with default parameters. To capture the primary sources of variability, the data was scaled, and 50 principal components (PCs) were computed from the scaled normalized counts of overdispersed genes via PCA analysis (using the RunPCA function with default settings). The number of PCs were selected for each sample individually. The nearest neighbors (using the FindNeighbors function with default parameters) was then identified and a Shared Nearest Neighbors (SNN) graph with a resolution of 0.8 (using the FindClusters function with default parameters) was constructed to detect cell clusters. Subsequently, the Seurat objects from each individual sample were combined using the merge function with default settings (SeuratObject, version 5.0.1). The combined data underwent normalization, identification of HVGs, and scaling. Following this, PCA analysis was conducted, nearest neighbors were identified, and finally the clusters were determined by generating an SNN graph with a resolution of 0.5, as described above. The result of analysis was then visualized using Uniform Manifold Approximation and Projection (UMAP) on the first 30 PCs with the RunUMAP function. Finally, the FindAllMarkers function was employed to identify differentially expressed genes (DEGs) within each cluster, using the following parameters: min.pct = 0.01, min.cells.feature = 3, min.cells.group = 3, logfc.threshold = 0.2, min.diff.pct = 0.2, and test.use = "wilcox". For downstream enrichment analysis, only DEGs with a p-value < 0.05 were selected. Additionally, after determining the identity of each cluster, the FindMarkers function was employed to identify DEGs between clusters from coculture and monoculture conditions at 8-week and 15-week time points, separately. The analysis was conducted using the following parameters: min.pct = 0.25, min.cells.feature = 3, min.cells.group = 3, logfc.threshold = 0.25, min.diff.pct = -Inf, and test.use = "wilcox".

### Gene ontology enrichment analysis

To verify the identity of each cluster, we performed Gene Ontology (GO) enrichment analysis on the list of upregulated genes (adjusted p-value < 0.05 and a log2 fold-change > 0.2) in each cluster using the Biological Process gene set library (p-values < 0.05). Additionally, cell type enrichment analysis was carried out using the CellMarker 2024 database (p-values < 0.05). All analyses were performed through the Enrichr database^17^ (https://maayanlab.cloud/Enrichr/).

### Integration analysis

To evaluate the similarity of cell clusters from the MyS and NMS model to their *in vivo* counterparts or to other established iPSC-derived *in vitro* models, we integrated our data with several developing^18,19^ and adult^20–22^ human brain single-nucleus/single-cell RNA sequencing (sn/scRNAseq) datasets as well as two *in vitro*^7,8^ models containing oligodendrocytes and OPCs. In the dataset from Fagiani *et al.*^8^, we specifically included samples that were treated with doxycycline but not with cerebrospinal fluid, encompassing both healthy and diseased samples. The integration of datasets was conducted using the merge function with default settings. The combined dataset was normalized, and HVGs were identified and scaled. PCA analysis was then performed, and nearest neighbors were identified based on the first 30 PCs. An SNN graph was constructed with a resolution of 0.5 to identify cell clusters. For batch correction, anchors between clusters were identified using the FindIntegrationAnchors function, employing the first 30 dimensions (Seurat package, version 5.0.3). These anchors were then used to integrate the datasets through IntegrateData function using default settings. The integrated data was normalized, and HVGs were identified and scaled. Next, the PCA analysis was done, clusters were identified and the SNN graph was generated as described above. Finally, the clusters were visualized using UMAP on the first 30 PCs with the RunUMAP function, as previously described. The resulting object was subsequently used for downstream analysis.

To assess the transcriptional similarity of oligodendrocyte populations across datasets, we performed a correlation analysis. As fetal brain datasets from 9-12 and 20-week-old fetuses lacked sufficient cells assignable to the oligodendrocyte cluster^18,19^, they were excluded from the analysis. For the remaining datasets, the AverageExpression function was used to calculate mean RNA expression values from the merged dataset to construct the correlation matrix. Top scoring features were subsequently selected using the SelectIntegrationFeatures function. The Spearman correlation was then determined on the processed data using the cor function. Finally, the correlation plot was generated utilizing the corrplot function from the corrplot package (version 0.95)^23^. To compare pseudobulk profiles of cell clusters from individual datasets within the merged dataset, the AggregateExpression function was employed (Seurat package, version 5.0.3). Subsequently, scatter plots were generated using the CellScatter function to visualize the expression levels of oligodendrocyte cell clusters across different datasets, with the myelin basic protein (MBP) specifically highlighted (Seurat package, version 5.0.3).

### SlingShot trajectory inference analysis

For trajectory inference, the batch-corrected dataset encompassing all datasets was utilized. The oligodendrocyte and fetal OPC clusters were subsetted using the Seurat package (version 5.0.3). Subsequently, trajectory analysis was conducted using the Slingshot package (version 2.12.0)^24^, with the OPC cluster from the youngest fetal sample^18^ designated as the root population.

### RNA velocity analysis

To investigate transcriptional dynamics and forecast cell fate decisions, we conducted RNA velocity analysis on a distinct differentiating path, including progenitor, proliferating OPC, OPC, and oligodendrocyte populations, for MyS model at 8- and 15-week time point. The RNA velocity analysis was executed using the Python package scVelo (0.3.2), following the guidelines established by Bergen *et al.* (2020)^25^ (https://scvelo.readthedocs.io/). Initially, clusters of interest were subsetted from the merged Seurat object of our snRNAseq data and converted to anndata format. The velocity command line tool was employed to calculate the spliced and unspliced counts matrix from the CellRanger output directory. Subsequently, the velocity was computed using the “dynamical mode”.

### Electrophysiology

MyS and NMS were harvested on week 8-10 and embedded in 2% low melting agarose (Sigma) diluted in PBS supplemented with 137 mM NaCl, 2.7 mM KCl, 10 mM Na_2_PO_4_, 1.8 mM KH_2_PO_4_, pH = 7.4. The embedding of the spheroids was done on ice and the solidified agarose containing spheroids were sectioned at a thickness of 300 μm, using Leica VT1200S vibratome in ice-cold oxygenated (95% O_2_ and 5% CO_2_) artificial cerebrospinal fluid solution, composed of 127 mM NaCl, 25 mM NaHCO_3_, 1.25 mM NaH_2_ PO_4_, 2.5 mM KCl, 1 mM MgCl_2_, 2 mM CaCl_2_, and 25 mM glucose. 2-3 sections from each spheroid (MyS or NM) were collected in 10 cm cell culture dish in NMM and the agarose was peeled off using a sharp blade under the microscope placed in a sterile culture hood. The spheroid slices of either MyS or NM were plated using a cut P1000 tip in one well each of high density multielectrode array (HD-MEA) 6 well plates (Maxwell Biosystems) coated with PLO-laminin. Excess NMM was carefully removed using SUGI absorption spears (AgnThos) while not disturbing the spheroid slices, and the slices were allowed to settle for a minute. 30 μL NMM supplemented with laminin was added very carefully, without disturbing the attachment of the spheroid slices in the well. These steps were repeated for all six wells and the HD-MEA plate was placed at 37 °C, 5% CO_2_ for 1-2 hours. Next, 150 μL medium was added to each well while avoiding disturbing the spheroid slices and the plates were returned to the incubator. The medium was changed on the following day with 150-200 mL OMM, followed by medium changes every other day. One day before the recording, OMM was replaced with BrainPhyS medium (Stemcell Technologies) and after the recording, BrainPhys was replaced with OMM. Spiking and bursting activity were recorded on a HD-MEA plate reader (Maxwell Biosystems) maintained at 37 °C and 5% CO_2_.

Whole-sample electrical recordings were obtained using the Activity Scan Assay. Recordings (MaxLab Live) were conducted for 30s per configuration. Spike times were extracted online using a threshold set at five standard deviations above the noise level. The high density of electrodes allowed precise identification and tracking of electrical activity in the MyS spheres for up to nine weeks post-plating. Wells were excluded from further analysis if threshold of 150 active electrodes with a firing rate (FR) of at least 1 Hz was not met, at any point during the recording period. The firing rate was calculated as the total number of spikes per electrode, normalized by the duration of the experiment. For burst detection, an inter-spike interval (ISI)-based method was used. Bursts were defined as sequences of at least four consecutive spikes with ISIs of less than 50 milliseconds. Additionally, the total number of active electrodes (defined as electrodes with a firing rate of at least 0.1 Hz) were calculated. Only active electrodes were used to compute the firing rate, inter-spike interval, burst rate, and inter-burst interval. All metrics were averaged across electrodes associated with each sphere half. Electrodes belonging to a sphere half were assigned based on the electrical map of firing rates generated at week 7, 8 or 9. If two distinct spheres were visually identifiable, active electrodes were manually assigned to each sphere and counted as two independent technical replicates. In cases where only one sphere was visible, all electrodes were attributed to that sphere. This method allowed us to distinguish the activity and maturation of individual sphere halves, as shown in Figure S8b. As the sphere halves exhibited distinct trends in electrophysiological activity over time, we further quantified these patterns by separating the number of spheres showing increased, decreased, or maintained activity levels. Activity between week 4 and 8 *in vitro* was compared, with spheres classified as ’increasing’ if they demonstrated at least a 30% rise in the number of active electrodes during this period, ‘decreasing’ if they showed a 30% decrease and ‘maintained’ otherwise.

### Lysolecithin treatment

Spheroids were incubated with lysolecithin (0.25 mg/mL, dissolved in OMM) for 15 hours in a 12 well plate on a shaker at 50 rpm, 37 °C and 5% CO_2_. Following lysolecithin treatment, spheroids were washed twice with PBS, resuspended in OMM, and returned to the shaker at 37 °C with 5% CO₂ for 3 hours. Spheroids were collected with cut P1000 tip and transferred to 96-well plate containing 4% PFA for overnight incubation at 4 °C. The samples were processed for immunofluorescence staining and TEM as described.

### RABV infection of MyS

Week 10 MyS were distributed in 96-well plates, with one spheroid per well. RABV mCherry-SAD-B19 or CVS-11 strains (30/100/300 TCID50 per spheroid) were added and incubated with the MyS for 24 hours. After incubation, the spheroids were washed 3 times with PBS, 300 µL fresh medium was added and the spheroids were further cultured. On day 1 and 4 post-infection, 150 µL cell culture supernatants from each well were collected for RT-qPCR analysis, and 200 µL fresh medium was added to each well to restore the volume. On day 6 post-infection, all culture medium were removed, spheroids were washed 3 times with PBS and fixed with 4% PFA at 4 °C overnight.

### Statistical analysis

The number of biological (N) and technical (n) replicates is mentioned in the respective figure legends. Summary of all the experiments with the number of replicates (N=biological, n=technical) and cell line used are mentioned in Table S8. Statistical analyses were performed using Prism v.8.2.1 (GraphPad). Tests used include two-way ANOVA with Sidak’s multiple comparisons test, one-way ANOVA with Sidak’s multiple comparisons test, paired and unpaired two-tailed t-tests. Differences were considered statistically significant for two-sided p-values < 0.05 (*p < 0.05; **p < 0.01; ***p < 0.001; ****p < 0.0001).

## Funding and Acknowledgements

This study was supported by the Research Foundation Flanders (FWO), including individual ongoing fellowships to K.A. (1S03424N) and T.B. (12AIK24N), and project grants to Y.C.C., C.V., K.A., R.R., F.N., J.T., E.E.C., K.N. (FWO-SBO-S001221N, OrganID and FWO-G0B5819N). Additional support was provided by the Mitialto Foundation. The authors gratefully acknowledge the VIB Bio Imaging Core (LIMONE) and KU Leuven Core facility financing for their support and assistance in this work. The computational resources and services used in this work were provided by the VSC (Flemish Supercomputer Center), funded by FWO and the Flemish Government.

## Contributions

KA designed and performed experiments, analyzed data and wrote the manuscript. RR, XW, CV, DJ, JN, YCC designed the experiments, helped with coordinating the study and contributed in manuscript editing. TB, GAm, KN, AS, GAr, EEC, FN, JT, KW, KV, NVD, SKP helped in the data analysis and edited the manuscript. IL, JS, LVDB, LDG, LM contributed in manuscript editing.

## Figures

**Figure S1:**
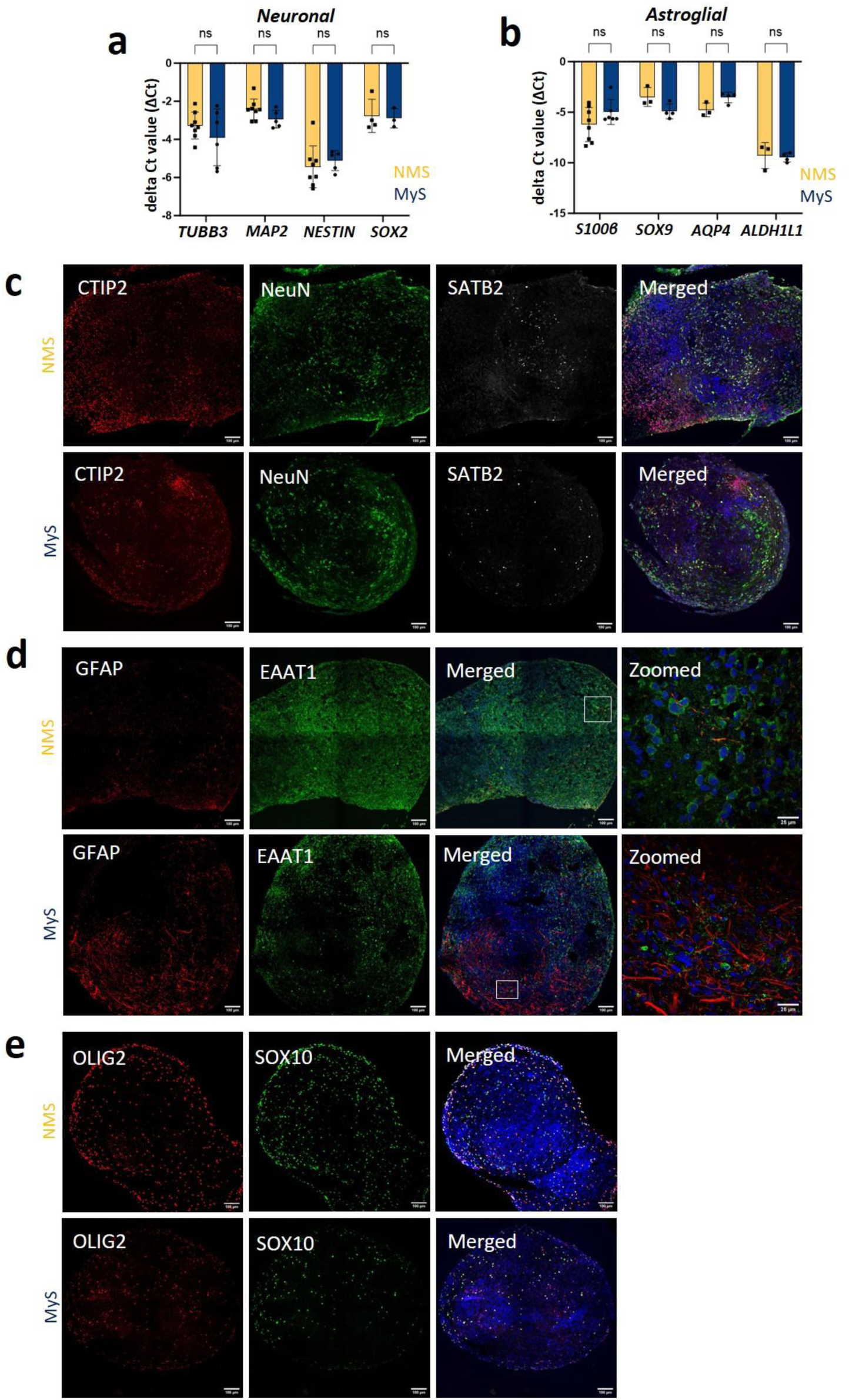
Cellular characterization of Myelin Spheres (MyS) and Neuron Monoculture Spheroids (NMS). **a-b** RT-qPCR of gene markers for neuronal (*TUBB3*, *MAP2*, *NESTIN*, *SOX2*) and astroglial (*S100β*, *SOX9*, *AQP4* and *ALDH1L1*) cells in week 10 MyS and NMS. **c-e** Immunostaining for neuronal (CTIP2, NeuN, SATB2), astroglia (GFAP, EAAT1) and oligodendroglia (OLIG2, SOX10) markers in week 10 MyS and NMS (scale bar = 100 μm and 25 μm for zoomed images (**d)**). Statistical analyses were performed by unpaired two tailed t-tests (**a-b**). Data shown are mean ± SD, N=3 (n=3-8, each data point represents two spheroids, **a-b**), representative immunostaining images from N=3 (**c-e**). ns = statistically not significant.

**Figure S2.**
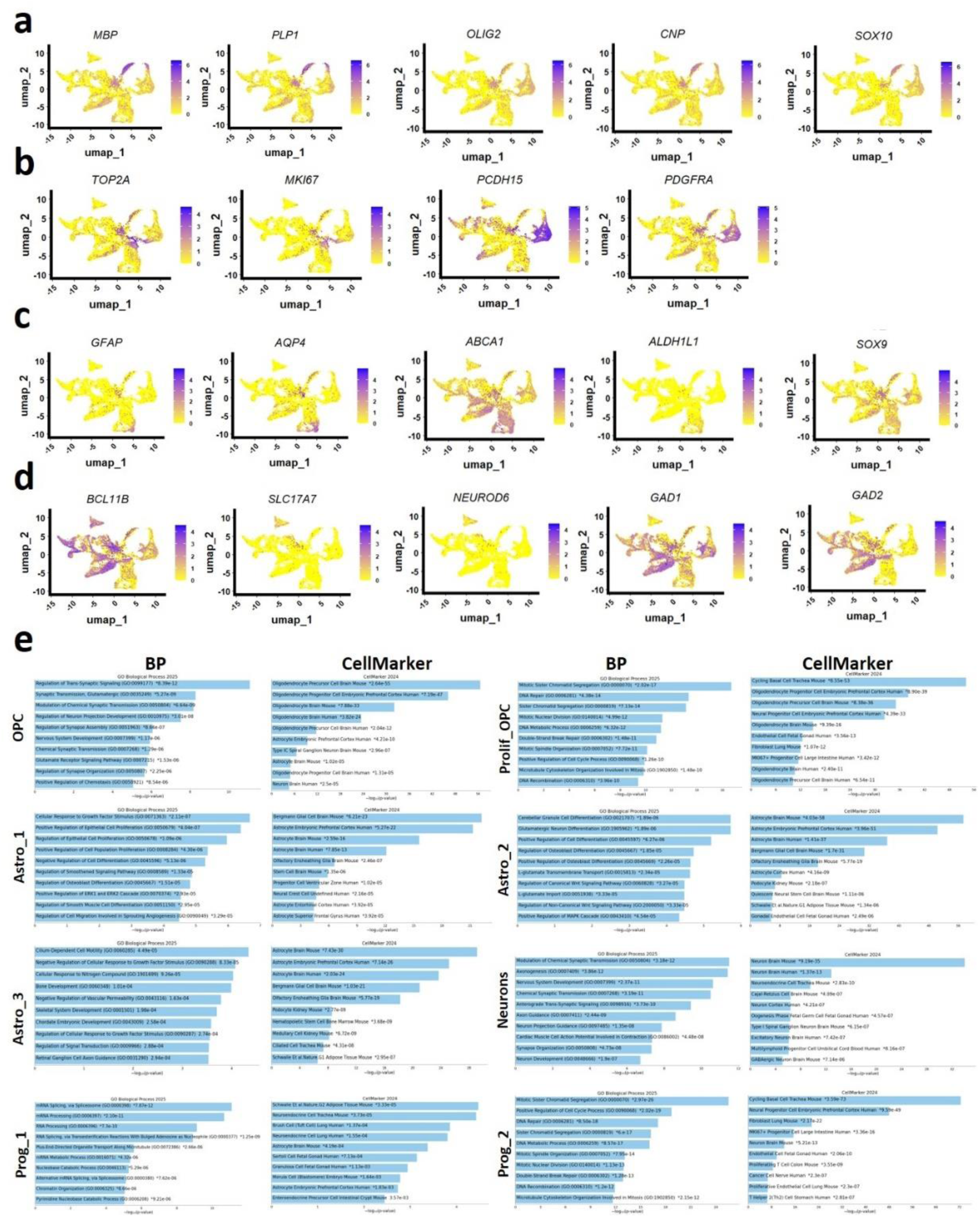
Characterization of the identities of cellular clusters in MyS and NMS. **a-d**: Feature plots illustrating marker gene expression for: **(a)** oligodendrocytes, **(b)** proliferating OPCs/OPCs, **(c)** astrocytes, and **(d)** neurons. **e** GO enrichment analysis for biological process terms and cell type enrichment analysis were conducted using the GO Biological Process 2025 and CellMarker 2024 databases. These analyses were based on (at least) top 500 DEGs of each cluster, with a p-adjusted value < 0.05 and an average log fold change > 0.20 for each cellular cluster. GO: Gene Ontology, BP: biological process, DEGs: differentially expressed genes.

**Figure S3.**
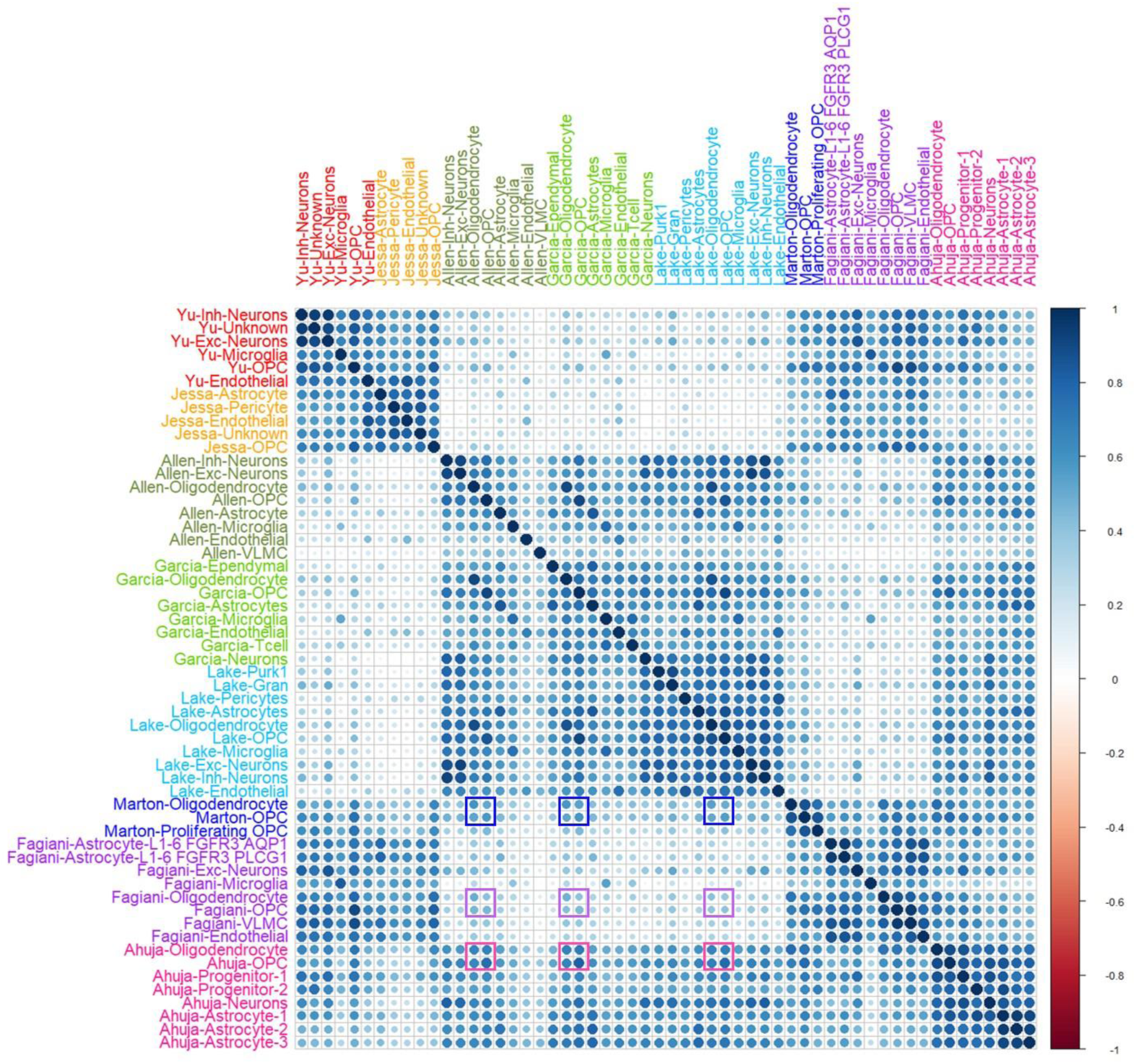
Analyses of transcriptional similarity between snRNAseq datasets from MyS and NMS, with sn/scRNAseq datasets from *in vivo* adult and developing brain cells as well as two *in vitro* brain models. The correlation plot demonstrates the correlations among all cellular clusters within the merged sn/scRNAseq datasets for adult^20–22^, fetal^18,19^ and *in vitro* datasets^7,8^. Correlations of Marton *et al.*^7^, Fagiani *et al.*^8^ and our dataset with adult brain shown in blue, violet and pink rectangles.

**Figure S4.**
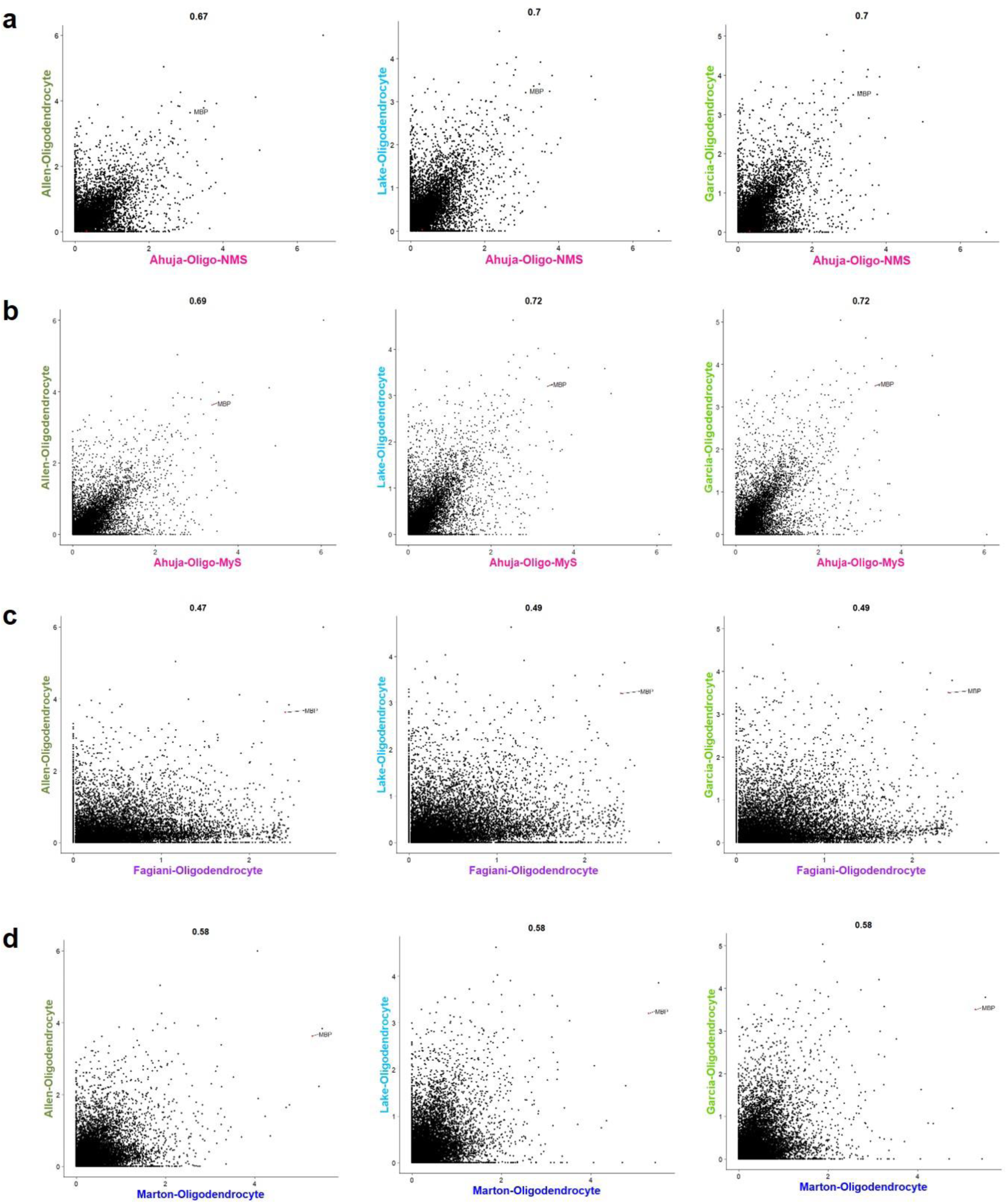
Comparative analysis of pseudobulk profiles of oligodendrocytes from MyS, NMS and *in vitro models* with the adult oligodendrocytes. **a-d:** The gene expression profiles of oligodendrocytes from (**a**) NMS, (**b**) MyS, (**c**) the Fagiani *et al.* model^8^, and (**d**) the Marton *et al.* model^7^ are compared to those of adult oligodendrocytes^20–22^.

**Figure S5.**
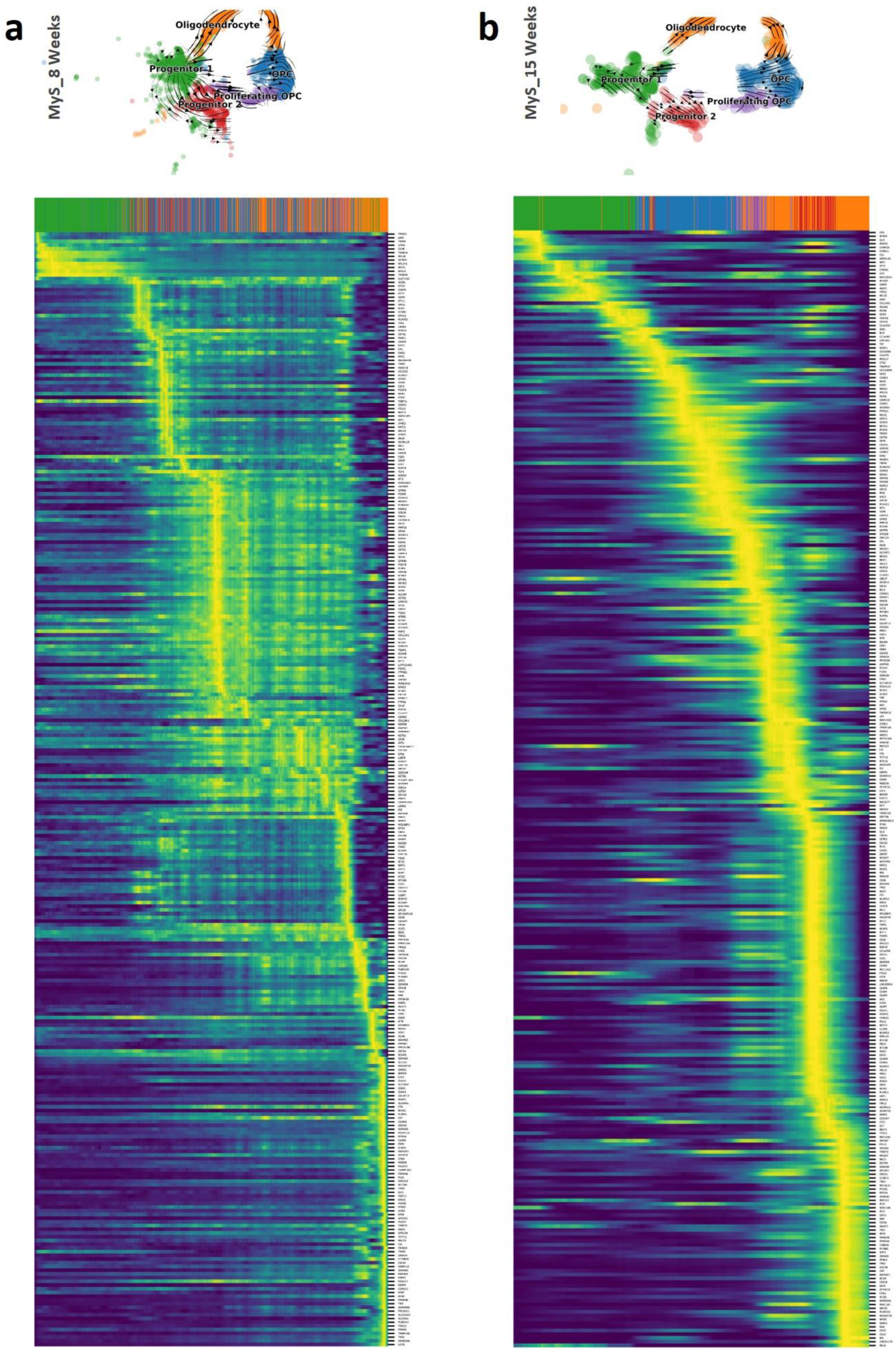
RNA velocity analysis demonstrated temporal dynamic of oligodendrocyte development in 8 and 15 Weeks MyS. **a-b:** The heatmaps revealed the dynamics of gene expression across latent-time of the underlying cellular processes, for the top 300 genes ranked by likelihood, in progenitors, OPCs, and oligodendrocyte clusters during (**a**) 8 weeks and (**b**) 15 weeks of culture.

**Figure S6:**
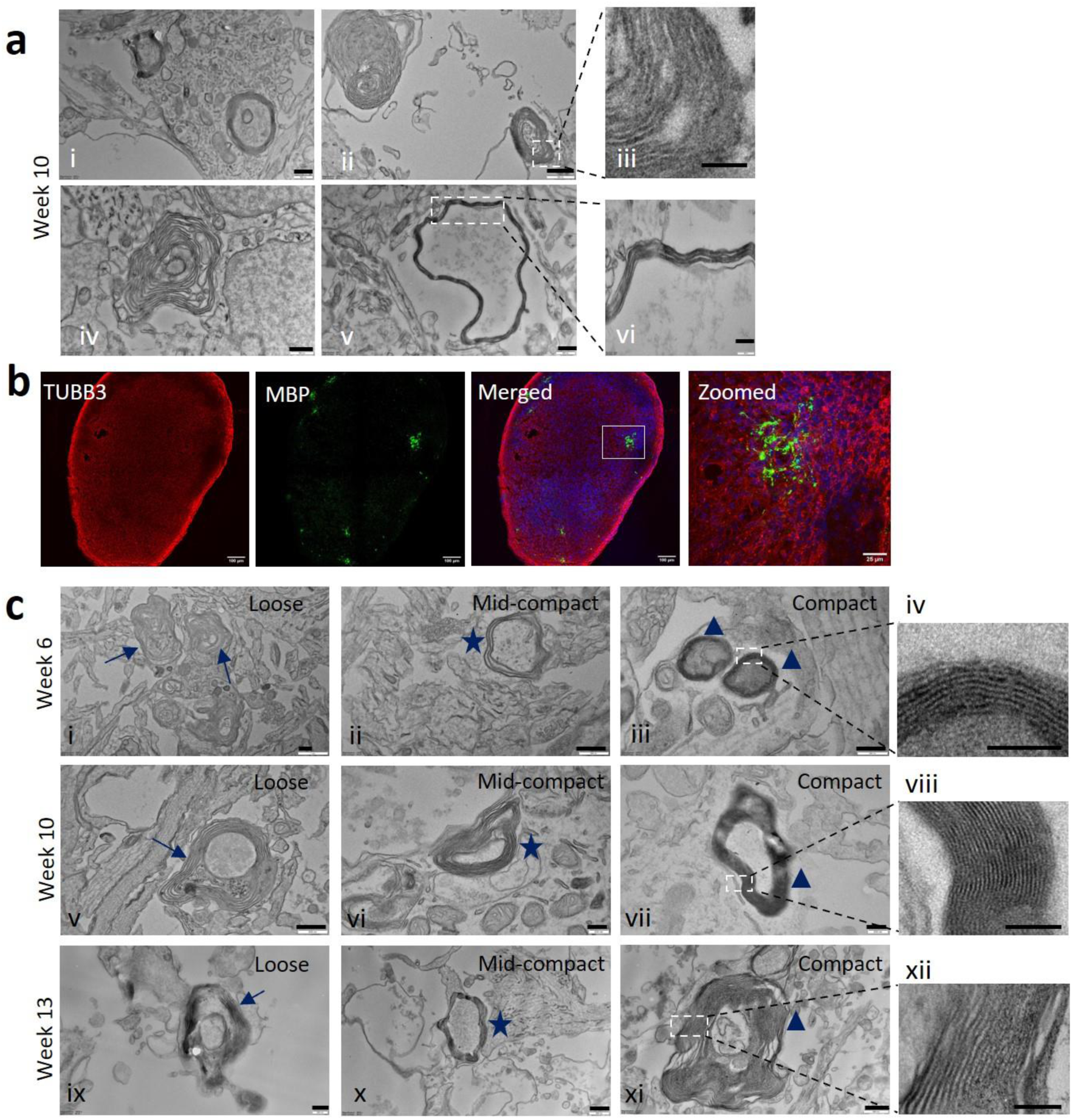
*In vitro* myelination in NMS generated from Sigma-iPSC0028 iPSCs and MyS-derived from human H9-ESCs. **a** TEM images showing formation of fewer myelin sheaths in week 10 NMS (scale bar = 500 nm (i, ii, iv, v) 100 nm (iii) and 200 nm (vi)). **b** Immunostaining for neuronal (TUBB3) and oligodendrocyte (MBP) markers in week 10 MyS-derived from H9-ESC cells (scale bar = 100 μm and 25μm for zoomed image). Nuclei were counterstained with Hoechst. **c** TEM images showing formation of different subtypes of myelin sheaths (loose (arrow) i, v, ix), mid-compact ((star) ii, vi, x) and compact ((triangle) iii, iv, vii, viii, xi, xii) in week 6, 10 and 13 H9-ESC derived MyS (scale bar = 500 nm (i, ii, v, x, xi), 200 nm (iii, vi, vii, ix), 100 nm (iv, viii), 250 nm (xii)). Representative images from N=3, n=3 (**a**), N=2, n=3 (**b-c**).

**Figure S7:**
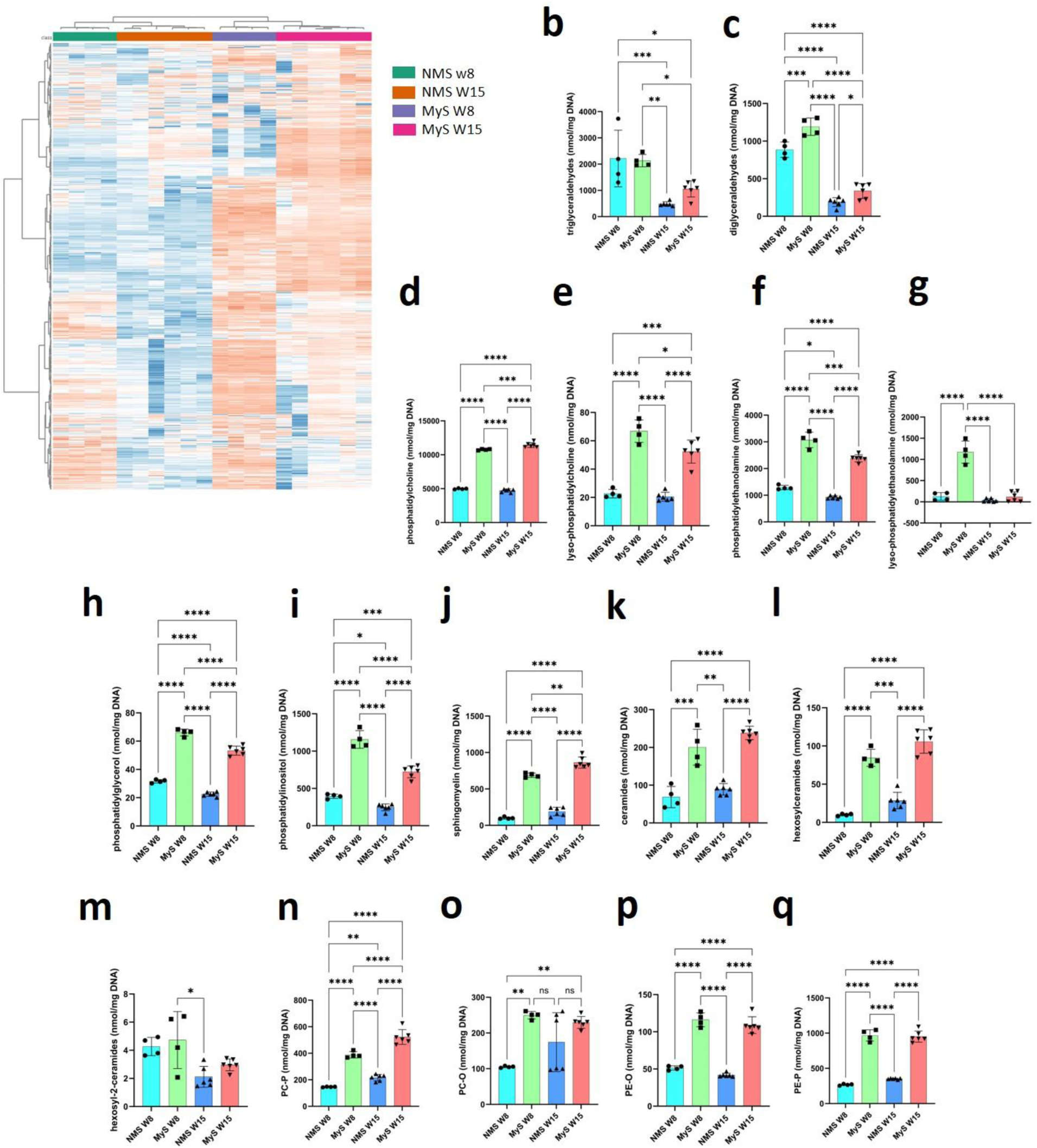
Global lipidome analysis revealed the presence of higher myelin lipids in MyS than NMS. **a** Heatmap showing separate clustering of MyS and NMS samples at week 8 and 15. **b-q** Absolute amounts of different subclasses of major lipid classes: triglycerides, diglycerides (glycerolipids, **b, c**), phosphatidylcholine (PC), lyso-PC, phosphatidylethanolamine (PE), lyso-PE, phosphatidylglycerol, phosphatidylinositol (glycerophospholipids, **d-i**), sphingomyelin, ceramides, hexosylceramides, hexosyl-2-ceramides (sphingolipids, **j-m**), PC-P, PC-O, PE-O and PE-P (esther linked phospholipids, **n-q**). Statistical analyses were performed by Two-way ANOVA with Sidak’s comparison test (**b-q**). Data shown are mean ± SD, N=4-6 (**b-q**). *p<0.05, **p<0.01, ***p<0.001, ****p<0.0001.

**Figure S8:**
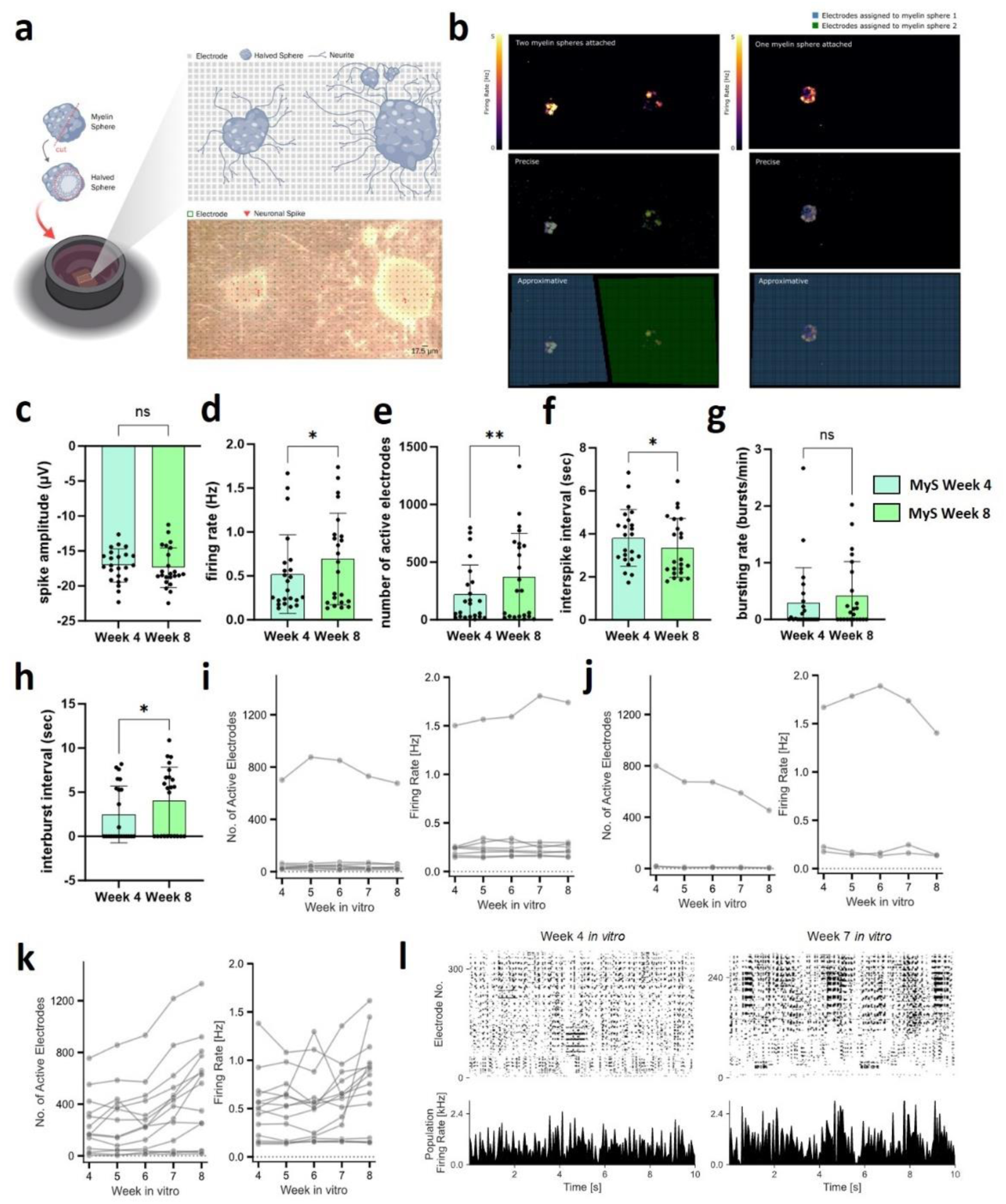
Electrophysiological characterization reveals functional activity in MyS. **a** Schematic illustration of MyS sliced spheres plated on a high density CMOS multi-electrode array for electrophysiological recordings. **b** Visualization of firing rate in separated electrode regions corresponding to individual MyS sliced spheres plated within a single well. **c-e** Quantification of neuronal spiking properties at week 4 and week 8, including spike amplitude (c), firing rate (d), number of active electrodes (e) and interspike interval (f) per well. **g-h** Assessment of neuronal bursting activity at week 4 and week 8, showing bursting rate (g) and interburst interval (h) per well. **i-k** Longitudinal analysis of individual MyS sliced spheres indicated by each graph lines, showing the number of active electrodes and the different firing rates over time: constant (i), decreasing (j), or increasing (k) activity patterns during weeks *in vitro*. **l** Representative raster plot illustrating increased spike activity in a MyS slice from week 4 to week 7 *in vitro*. Statistical analyses were performed by paired two tailed *t*-tests (**c-h**). Data shown are mean ± SD for N = 4 independent experiments, n = 23 MyS sliced spheres, *p< 0.05, **p< 0.01. ns = not significant.

**Figure S9:**
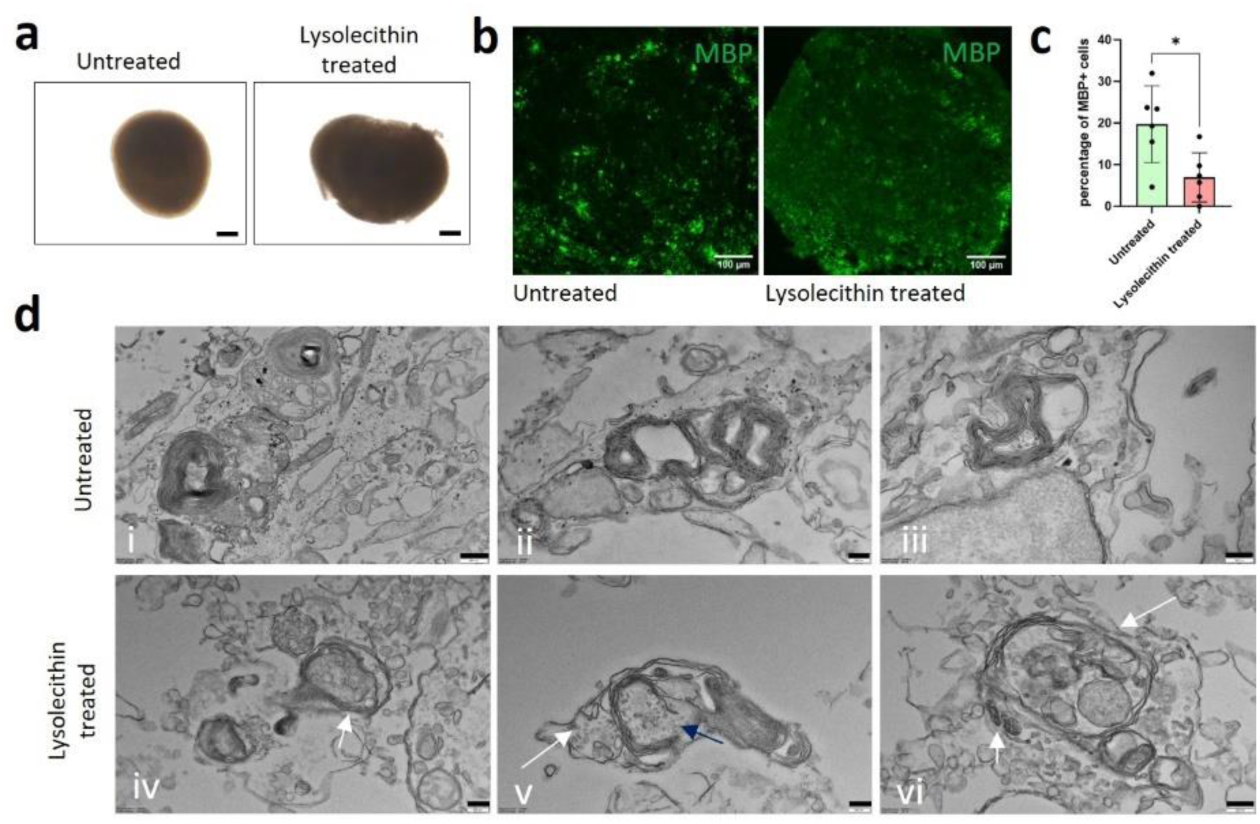
Lysolecitin induced demyelination in MyS. **a** Brightfield image of untreated and lysolecithin-treated week 10 MyS (scale bar = 200 μm). **b** Immunostaining of mature oligodendrocyte marker (MBP) in untreated and lysolecithin-treated week 10 MyS (scale bar = 100 μm). **c** Quantification of immunostained MBP^+^ cells in untreated and lysolecithin-treated week 10 MyS. **d** TEM images showing lysed myelin sheaths (arrows) in lysolecithin-treated week 10 MyS as compared to untreated week 10 MyS (scale bar = 500 nm-i; 200nm-ii-vi). Statistical analyses were performed by unpaired two tailed *t*-tests (**c**). Data shown are ± SD, representative images and quantification from N = 3 (n = 5-6, each data point indicates 3 slices) (**a-d**). *p<0.05.

**Figure S10:**
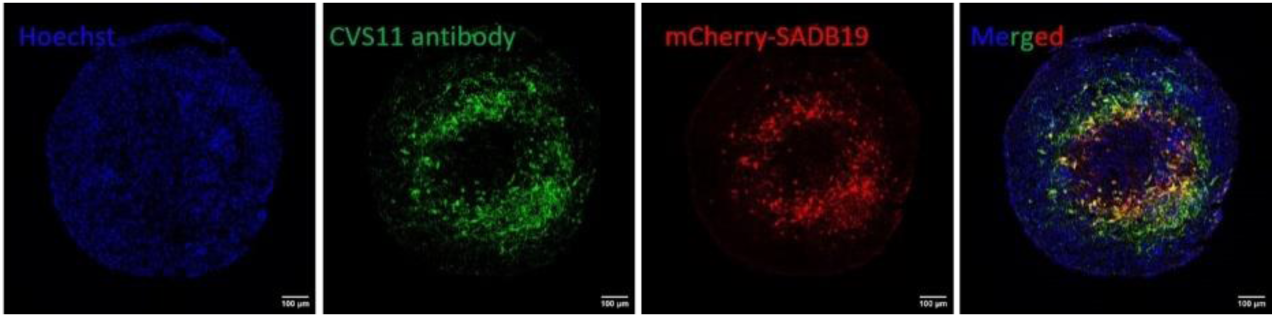
mCherry-SAD-B19 signal corresponds to RABV particles in MyS. Week 10 MyS infected with mCherry-SAD-B19 stained with an anti-RABV N protein antibody against the RABV Nucleoprotein shows colocalization of mCherry signal with RABV infected cells (scale bar = 100 μm).

**Figure S11:**
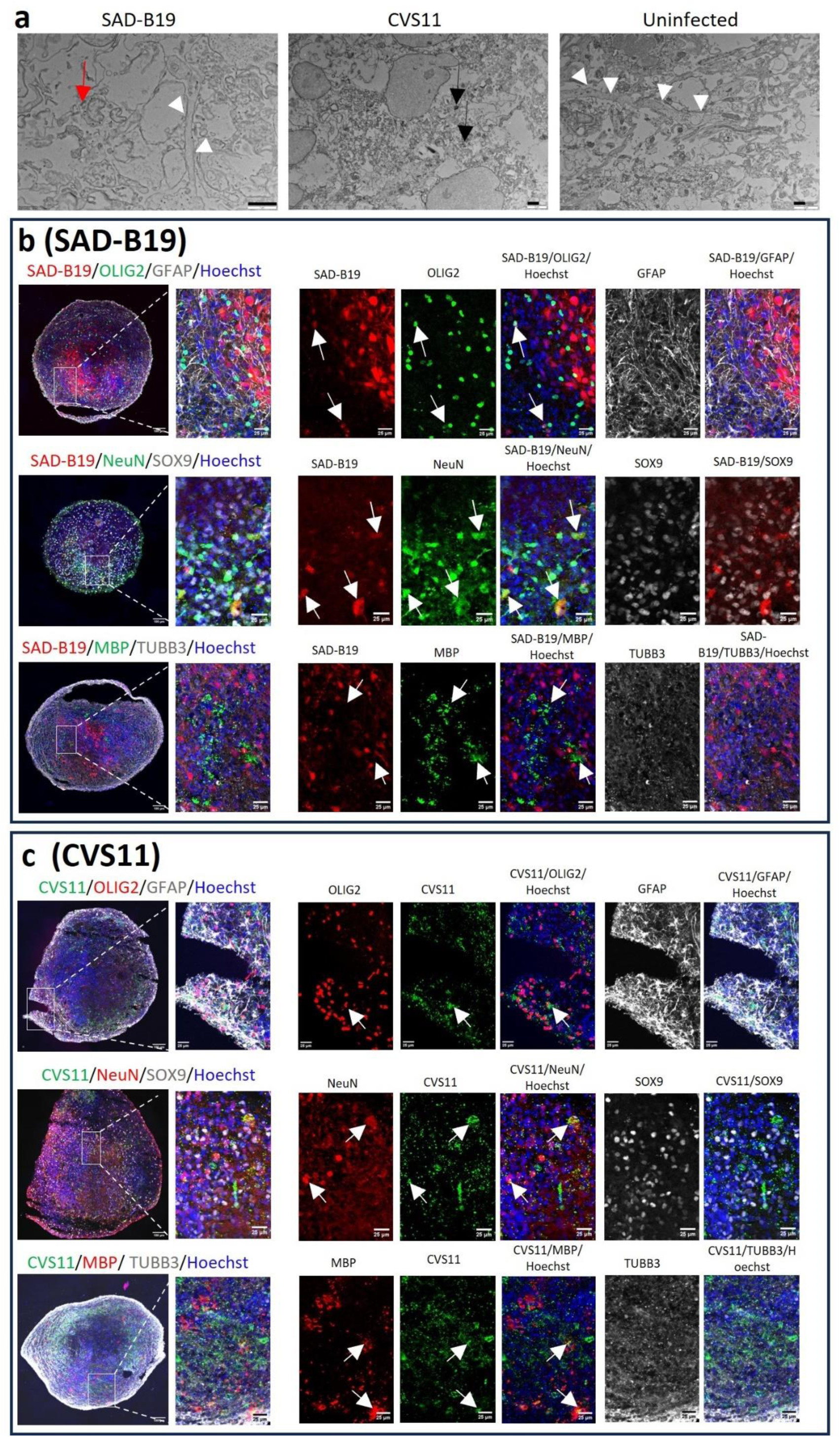
Differences in Rabies virus pathology and tropism observed across strains in MyS. **a** Transmission electron microscopic images showing severe damage to microtubule networks (white arrow heads) by the CVS-11 (black arrow) but not mCherry-SAD-B19 (red arrow) virus as compared to uninfected week 10 MyS (scale bar = 1μm). **b** Immunostaining imaging for markers of neuron (NeuN, TUBB3), astroglia (GFAP, SOX9), oligodendrocyte progenitors (OLIG2) and mature oligodendrocytes (MBP) along with mCherry expression from mCherry-SAD-B19 virus demonstrating tropism towards neurons, and oligodendrocyte progenitors but not astroglia and mature oligodendrocytes. **c** Immunostaining imaging for neurons (NeuN, TUBB3), astroglia (GFAP/SOX9), oligodendrocyte progenitors (OLIG2) and mature oligodendrocytes (MBP) combined with anti-RABV Nucleoprotein antibody demonstrating tropism of the CVS-11 virus towards neurons, oligodendrocyte progenitors and mature oligodendrocytes but not to astrocytes (scale bar = 100 μm and 25μm for the zoomed panels). Zoomed panels are cropped manually from their corresponding images for better visualization. Representative images from N=3 (**a-c**).

**Video S1: 3D reconstruction of TEM image showing myelination in Sigma iPSC-derived MyS.** Scale bar = 500nm.

**Video S2: 3D reconstruction of TEM image showing myelination in H9-ESC derived MyS.** Scale bar = 500nm.

## Bibliography

1. Gargareta, V. I. et al. Conservation and divergence of myelin proteome and oligodendrocyte transcriptome profiles between humans and mice. Elife 11, (2022).

2. Chrast, R., Saher, G., Nave, K. A. & Verheijen, M. H. G. Lipid metabolism in myelinating glial cells: lessons from human inherited disorders and mouse models. J. Lipid Res. 52, 419–434 (2011).

3. Piekarski, D. J. et al. White matter microstructural integrity continues to develop from adolescence to young adulthood in mice and humans: Same phenotype, different mechanism. Neuroimage: Reports 3, 100179 (2023).

4. Madhavan, M. et al. Induction of myelinating oligodendrocytes in human cortical spheroids. Nat. Methods 15, 700–706 (2018).

5. Feng, L., et al. Developing a human iPSC-derived three-dimensional myelin spheroid platform for modeling myelin diseases. iScience 26, (2023).

6. van Tilborg, E. et al. Origin and dynamics of oligodendrocytes in the developing brain: Implications for perinatal white matter injury. Glia 66, 221 (2017).

7. Marton, R. M. et al. Differentiation and Maturation of Oligodendrocytes in Human Three-Dimensional Neural Cultures. Nat. Neurosci. 22, (2019).

8. Fagiani, F. et al. A glia-enriched stem cell 3D model of the human brain mimics the glial-immune neurodegenerative phenotypes of multiple sclerosis. Cell Reports Med. 5, (2024).

9. Shi, Y., Kirwan, P. & Livesey, F. J. Directed differentiation of human pluripotent stem cells to cerebral cortex neurons and neural networks. Nat. Protoc. 2012 710 7, 1836–1846 (2012).

10. García-León, J. A. et al. Generation of oligodendrocytes and establishment of an all-human myelinating platform from human pluripotent stem cells. Nat. Protoc. 15, 3716– 3744 (2020).

11. Morriss, N. J. et al. Automated Quantification of Immunohistochemical Staining of Large Animal Brain Tissue Using QuPath Software. Neuroscience 429, 235–244 (2020).

12. Wu, G. et al. A simplified method for measuring neutralising antibodies against rabies virus. J. Virol. Methods 319, (2023).

13. Peng, H., Bria, A., Zhou, Z., Iannello, G. & Long, F. Extensible visualization and analysis for multidimensional images using Vaa3D. Nat. Protoc. 2013 91 9, 193–208 (2014).

14. Pang, Z. et al. MetaboAnalyst 6.0: towards a unified platform for metabolomics data processing, analysis and interpretation. Nucleic Acids Res. 52, W398–W406 (2024).

15. Ewald, J. D. et al. Web-based multi-omics integration using the Analyst software suite. Nat. Protoc. 19, 1467–1497 (2024).

16. Karpievitch, Y. V., Nikolic, S. B., Wilson, R., Sharman, J. E. & Edwards, L. M. Metabolomics Data Normalization with EigeNMS. PLoS One 9, e116221 (2014).

17. Chen, E. Y. et al. Enrichr: Interactive and collaborative HTML5 gene list enrichment analysis tool. BMC Bioinformatics 14, 1–14 (2013).

18. Yu, Y. et al. Interneuron origin and molecular diversity in the human fetal brain. Nat. Neurosci. 2021 2412 24, 1745–1756 (2021).

19. Jessa, S. et al. Stalled developmental programs at the root of pediatric brain tumors. Nat. Genet. 2019 5112 51, 1702–1713 (2019).

20. Bakken, T. E. et al. Comparative cellular analysis of motor cortex in human, marmoset and mouse. Nat. 2021 5987879 598, 111–119 (2021).

21. Lake, B. B. et al. Integrative single-cell analysis of transcriptional and epigenetic states in the human adult brain. Nat. Biotechnol. 2017 361 36, 70–80 (2017).

22. Garcia, F. J. et al. Single-cell dissection of the human brain vasculature. Nat. 2022 6037903 603, 893–899 (2022).

23. Friendly, M. Corrgrams. Am. Stat. 56, 316–324 (2002).

24. Street, K. et al. Slingshot: Cell lineage and pseudotime inference for single-cell transcriptomics. BMC Genomics 19, 1–16 (2018).

25. Bergen, V., Lange, M., Peidli, S., Wolf, F. A. & Theis, F. J. Generalizing RNA velocity to transient cell states through dynamical modeling. Nat. Biotechnol. 2020 3812 38, 1408–1414 (2020).

26. Miyazaki, Y. et al. Oligodendrocyte-derived LGI3 and its receptor ADAM23 organize juxtaparanodal Kv1 channel clustering for short-term synaptic plasticity. Cell Rep. 43, 113634 (2024).

27. Zhou, L. et al. Gab1 mediates PDGF signaling and is essential to oligodendrocyte differentiation and CNS myelination. Elife 9, (2020).

28. Santra, M. et al. Thymosin β4 Up-regulation of MicroRNA-146a Promotes Oligodendrocyte Differentiation and Suppression of the Toll-like Proinflammatory Pathway. J. Biol. Chem. 289, 19508 (2014).

29. Wang, P. et al. Predicting signaling pathways regulating demyelination in a rat model of lithium-pilocarpine-induced acute epilepsy: A proteomics study. Int. J. Biol. Macromol. 193, 1457–1470 (2021).

30. Chapman, T. W., Piedra, E. T. & Hill, R. A. Oligodendrocyte maturation alters the cell death mechanisms that cause demyelination. doi:10.1101/2023.09.26.557781.

31. Chamling, X. et al. Single-Cell Transcriptomic Analysis Reveals Molecular Diversity of Human Oligodendrocyte Progenitor Cells. bioRxiv 2020.10.07.328971 (2020) doi:10.1101/2020.10.07.328971.

32. Ji, K., Ren, H., Zhao, X. & Yan, C. Migratory Rolandic Encephalopathy Caused by the Mitochondrial ND3 Variant. Neurology 98, 80–81 (2022).

33. Hassel, L. A. et al. Differential activity of transcription factor Sox9 in early and adult oligodendroglial progenitor cells. Glia 71, 1890–1905 (2023).

34. Braccioli, L., Vervoort, S. J., Puma, G., Nijboer, C. H. & Coffer, P. J. SOX4 inhibits oligodendrocyte differentiation of embryonic neural stem cells in vitro by inducing Hes5 expression. Stem Cell Res. 33, 110–119 (2018).

35. Wang, Y. et al. PARP1-mediated PARylation activity is essential for oligodendroglial differentiation and CNS myelination. Cell Rep. 37, (2021).

36. Yaffe, Y. et al. The myelin proteolipid plasmolipin forms oligomers and induces liquid-ordered membranes in the Golgi complex. J. Cell Sci. 128, 2293–2302 (2015).

37. Reza, S., Ugorski, M. & Suchański, J. Glucosylceramide and galactosylceramide, small glycosphingolipids with significant impact on health and disease. Glycobiology 31, 1416 (2021).

38. Huang, H. et al. Tmeff2 is expressed in differentiating oligodendrocytes but dispensable for their differentiation in vivo. Sci. Reports 2017 71 7, 1–8 (2017).

39. Filippini, A. et al. Leucine-Rich Repeat Kinase-2 Controls the Differentiation and Maturation of Oligodendrocytes in Mice and Zebrafish. Biomolecules 14, (2024).

40. Lin, J. P., Mironova, Y. A., Shrager, P. & Giger, R. J. LRP1 regulates peroxisome biogenesis and cholesterol homeostasis in oligodendrocytes and is required for proper CNS myelin development and repair. Elife 6, e30498 (2017).

41. Bergen, V., Lange, M., Peidli, S., Wolf, F. A. & Theis, F. J. Generalizing RNA velocity to transient cell states through dynamical modeling. Nat. Biotechnol. 2020 3812 38, 1408–1414 (2020).

42. Vana, N. S. et al. Early cortical oligodendrocyte precursor cells are transcriptionally distinct and lack synaptic connections. Glia 71, 2210–2233 (2023).

43. Khawaja, R. R. et al. GluA2 overexpression in oligodendrocyte progenitors promotes postinjury oligodendrocyte regeneration. Cell Rep. 35, 109147 (2021).

44. Fard, M. K. et al. BCAS1 expression defines a population of early myelinating oligodendrocytes in multiple sclerosis lesions. Sci. Transl. Med. 9, (2017).

45. Kim, D., An, H., Fan, C. & Park, Y. Identifying oligodendrocyte enhancers governing Plp1 expression. Hum. Mol. Genet. 30, 2225–2239 (2021).

46. Fulton, D., Paez, P. M. & Campagnoni, A. T. The multiple roles of myelin protein genes during the development of the oligodendrocyte. ASN Neuro 2, e00027 (2010).

47. Kaneko, N. et al. ADAMTS2 promotes radial migration by activating TGF-β signaling in the developing neocortex. EMBO Rep. 25, 3090–3115 (2024).

48. Fitzner, D. et al. Cell-Type- and Brain-Region-Resolved Mouse Brain Lipidome. Cell Rep. 32, 108132 (2020).

49. Saher, G. et al. High cholesterol level is essential for myelin membrane growth. Nat. Neurosci. 2005 84 8, 468–475 (2005).

50. Berghoff, S. A. et al. Neuronal cholesterol synthesis is essential for repair of chronically demyelinated lesions in mice. Cell Rep. 37, 109889 (2021).

51. Birgbauer, E., Rao, T. S. & Webb, M. Lysolecithin induces demyelination in vitro in a cerebellar slice culture system. J. Neurosci. Res. 78, 157–166 (2004).

52. Morecki, R. & Zimmerman, H. M. Human Rabies Encephalitis: Fine Structure Study of Cytoplasmic Inclusions. Arch. Neurol. 20, 599–604 (1969).

53. Matsumoto, S. ELECTRON MICROSCOPE STUDIES OF RABIES VIRUS IN MOUSE BRAIN. J. Cell Biol. 19, 565–591 (1963).

54. Harsha, P. K. et al. Mitochondrial Dysfunction in Rabies Virus-Infected Human and Canine Brains. Neurochem. Res. 47, 1610–1636 (2022).

55. Li, X.-Q., Sarmento, L. & Fu, Z. F. Degeneration of Neuronal Processes after Infection with Pathogenic, but Not Attenuated, Rabies Viruses. J. Virol. 79, 10063–10068 (2005).

56. Giussani, P., Prinetti, A. & Tringali, C. The role of Sphingolipids in myelination and myelin stability and their involvement in childhood and adult demyelinating disorders. J. Neurochem. 156, 403–414 (2021).

57. Stikov, N. et al. In vivo histology of the myelin g-ratio with magnetic resonance imaging. Neuroimage 118, 397–405 (2015).

58. Bouhrara, M. et al. Age-related estimates of aggregate g-ratio of white matter structures assessed using quantitative magnetic resonance neuroimaging. Hum. Brain Mapp. 42, 2362–2373 (2021).

59. Peters, A., Palay, S. L. & Webster, H. deF. The Fine Structure of the Nervous System: neurons and their supporting cells. xviii, 494 (1976).

60. Tsiang, H., Koulakoff, A., Bizzini, B. & Berwald-Netter, Y. Neurotropism of rabies virus. An in vitro study. J. Neuropathol. Exp. Neurol. 42, 439–452 (1983).

61. Potratz, M. et al. Astrocyte Infection during Rabies Encephalitis Depends on the Virus Strain and Infection Route as Demonstrated by Novel Quantitative 3D Analysis of Cell Tropism. Cells 9, 412 (2020).

62. Potratz, M. et al. Neuroglia infection by rabies virus after anterograde virus spread in peripheral neurons. Acta Neuropathol. Commun. 8, 199 (2020).

63. Jenson, A. B., Rabin, E. R., Bentinck, D. C. & Melnick, J. L. Rabiesvirus neuronitis. J. Virol. 3, 265–269 (1969).

64. Murphy, F. A., Bauer, S. P., Harrison, A. K. & Winn, W. C. Comparative pathogenesis of rabies and rabies-like viruses. Viral infection and transit from inoculation site to the central nervous system. Lab. Investig. 28, 361–376 (1973).

65. Mancuso, R. et al. Xenografted human microglia display diverse transcriptomic states in response to Alzheimer’ s disease-related amyloid-β pathology. Nat. Neurosci. 2024 275 27, 886–900 (2024).

66. Garton, T., Gadani, S. P., Gill, A. J. & Calabresi, P. A. Neurodegeneration and demyelination in multiple sclerosis. Neuron 112, 3231–3251 (2024).

